# In-Scanner Thoughts shape Resting-state Functional Connectivity: how participants “rest” matters

**DOI:** 10.1101/2024.06.05.596482

**Authors:** J Gonzalez-Castillo, MA Spurney, KC Lam, IS Gephart, F Pereira, DA Handwerker, JWY Kam, PA Bandettini

## Abstract

Resting-state fMRI (rs-fMRI) scans—namely those lacking experimentally-controlled stimuli or cognitive demands—are often used to identify aberrant patterns of functional connectivity (FC) in clinical populations. To minimize interpretational uncertainty, researchers control for across-cohort disparities in age, gender, co-morbidities, and head motion. Yet, studies rarely, if ever, consider the possibility that systematic differences in inner experience (i.e., what subjects think and feel during the scan) may directly affect FC measures. Here we demonstrate that is the case using a rs-fMRI dataset comprising 471 scans annotated with experiential data. Wide-spread significant differences in FC are observed between scans that systematically differ in terms of reported in-scanner experience. Additionally, we show that FC can successfully predict specific aspects of in-scanner experience in a manner similar to how it predicts demographics, cognitive abilities, clinical outcomes and labels. Together, these results highlight the key role of in-scanner experience in shaping rs-fMRI estimates of FC.

## Introduction

Resting-state scans are a central component of large-scale, international neuroimaging efforts aimed at mapping the human functional connectome in-vivo, in both healthy and patient populations^1^. This prominence of resting-state in cognitive and clinical neuroscience research is motivated by its experimental simplicity, short scanning time (which translates into cost effectiveness), richness of information, and low demands on participants (typical instructions might be "*remain still and awake as you let your mind wander freely")*. Yet, despite its wide adoption, we still lack an understanding of the nature and biological purpose of the neural processes that give rise to the observed synchronized patterns of spontaneous activity during rest^2,3^. This knowledge gap severely limits both our ability to interpret results and to develop reliable brain-based biomarkers of disease.

Neural mechanisms previously suggested as explanatory phenomena for resting-state functional connectivity patterns (rsFC) include memory consolidation^3^, learning^4,5^, fluctuations in wakefulness^6,7^, homeostatic processes needed to maintain the brain’s functional integrity^3^, the replay and update of predictive models of the environment^8^, processing of interoceptive signals from the rest of the body^9^, and those associated with different aspects of the subjective experience participants undergo while being scanned^2^. Of these, the role of the in-scanner subjective experience is perhaps the least established^10^. This is because researchers rarely collect information about what participants think and feel inside the scanner. A few instances where such data is available suggest that some aspects of in-scanner experience can be related to resting-state signals. For example, reported levels of imagery correlate with the strength of resting-state fluctuations in perilingual cingulate cortex^11,12^, and reported levels of comfort with that of somatosensory cortex^12^; a higher propensity to focus in-scanner thoughts on current concerns (as opposed to future planning) has been linked to stronger betweenness centrality for the left middle frontal gyrus^13^ (a key component of the salience network); and the qualia of inner voices (i.e., inner hearing vs. inner speaking) was associated with differential patterns of spontaneous activity during rest in language related areas^14^.

Most of these reports focus on one aspect of internal thoughts without systematically modeling how a broader range of in-scanner experiences might influence rsFC patterns. This matters because unmodeled variance based on these differences could hide aberrant patterns of inter-regional communication linked to clinical symptoms and hinder efforts aimed at developing brain-based biomarkers of disease progression and therapeutic efficacy. In fact, some have argued for the use of naturalistic experiment designs in detriment of resting-state, due to the many interpretational challenges stemming from resting-state lacking a well-defined relationship with cognitive processes^15^. Conversely, others have argued that state-level factors, such as variations in in-scanner experience, might have a minimal or no modulatory effect at all on rsFC^3^. To shed light on these debates, which are key for forecasting the true clinical potential of rs-fMRI (and thus for making funding decisions), and understand the role that in-scanner experience might play in common research applications of rsfMRI, we seek to answer three specific questions.

First, are patterns of in-scanner thoughts during rest characteristic and consistent in individuals over time? If so, putative modulatory effects of in-scanner experience in rsFC will show some level of stability across repeated scans, potentially confounding (or explaining) the relationship between rsFC and other subject-specific, trait-level characteristics of interest. It would also mean that such effects could not be easily removed by simple averaging across successive scanning sessions.

Second, do scans that systematically differ in terms of thought content show significant differences in rsFC? If so, are such differences interpretable and in agreement with our current understanding of the functional organization of the brain? A common way to uncover systems-level FC correlates of brain disease is to acquire resting-state scans in two groups—those afflicted by the condition of interest and healthy controls—and to contrast those after careful pre-processing and matching of samples in terms of head motion and basic demographics. Observed differences that survive strict statistical testing are then interpreted as being related to the symptomatology being examined^16^. Rarely do researchers consider the possibility that such observations, in full or partially, might be explained by systematic differences in in-scanner experience across the groups. To evaluate whether this should be a concern, we take hundreds of scans from a healthy population and subdivide them into sets that differ in terms of thought content, but are otherwise similar regarding sample size, gender, age, head motion, and wakefulness. Should a comparison across these sets render widespread significant differences, this will suggest that in-scanner experience should be considered a potential explanatory factor in rsFC studies of populations known to have altered spontaneous thought patterns, such as it is the case for those afflicted by dysphoria^17^, major depressive disorder^18,19^, autism spectrum disorder^20^ and borderline personality disorder^21^; to name a few.

Third, is it possible to predict different aspects of in-scanner thoughts using rsFC, in a manner similar to how other studies predict phenotypical scores related to personality traits^22^, cognitive abilities^23,24^ and clinical symptoms or labels^25,26^? If so, this would suggest that the imprints of in-scanner experience in rsFC are equivalently strong to those of these other phenotypical characteristics commonly targeted by resting-state research; and can be potential targets for better interpretation of clinically-relevant variations.

Here, we answer these three questions affirmatively, using a publicly available sample of 471 resting-state scans^27^. These scans are annotated with retrospective debriefing self-reports about in-scanner thoughts and wakefulness levels (Table 1). These annotations, although limited in scope, provide unique and valuable information about what participants experienced while being scanned.

**Table 1.** Experiential Questions following the completion of each resting-state scan.

| TARGET | LABEL | STATEMENT |
| --- | --- | --- |
| CONTENT OF THOUGHTS | Surroundings | I thought about my environment/surroundings |
|  | People | I thought about other people |
|  | Myself | I thought about myself |
|  | Past | I thought about past events |
|  | Future | I thought about future events |
|  | Negative | I thought about something negative |
|  | Positive | I thought about something positive |
| FORM OF THOUGHTS | Intrusive | My thoughts were intrusive |
|  | Specific | My thoughts were more specific than vague |
|  | Words | My thoughts were in the form of words |
|  | Images | My thoughts were in the form of images |
| WAKEFULNESS | Wakefulness | I was completely awake |

Our results highlight the important role of in-scanner experience as a key factor driving rsfMRI measurements. They also suggest that putative systematic differences in in-scanner thought patterns (e.g., excessive ruminative thoughts, disparate focus of attention, etc.) across surveyed populations can cloud the interpretation of population studies by adding one more source of unmodeled variance. To mitigate these issues, and facilitate more detailed interpretations of results, future resting-state studies should consider the inclusion of experiential self-reports as a companion to every scan. This is a relatively simple addition, compared to the cost and complexity of executing a single resting-state scan, and would enrich resting-state datasets with new opportunities for analysis and discovery. For example, it would allow the modeling of in-scanner thoughts as additional confounds (in a manner similar to head motion or demographics) when such mental aspects are not the target of inquiry. This could, in turn, improve the accuracy of existing rsfMRI predictive models and uncover new brain-behavior relationships. Similarly, such richer resting-state datasets would also allow the investigation of the neural correlates of self-generated thought processes such as creative thinking^28^, mind-wandering^29^, interoceptive awareness^30^, intrusive thinking^31^ and mind blanking^32^.

## Results

### Self-report annotations of resting-state scans

At the end of each resting-state scan, participants filled out the short New York City Questionnaire (sNYCQ, Table 1) that included seven statements about the content of their thoughts during the scan (i.e., “*I thought about my environment/surroundings*”), four about the form of the thoughts (i.e., “*My thoughts were in the form of images*”), and one about the subjectively perceived level of wakefulness (i.e., “*I was completely awake*”). For each statement, participants responded with a number between 0 (“Describes my thoughts not at all”) and 100 (“Describes my thoughts completely”) (Fig. 1a). Because the modulatory role of wakefulness on rsFC is already well-stablished^6,7,33^, we kept the answers to the wakefulness statement separate to the rest of the questionnaire during the analyses.

**Figure. 1.**
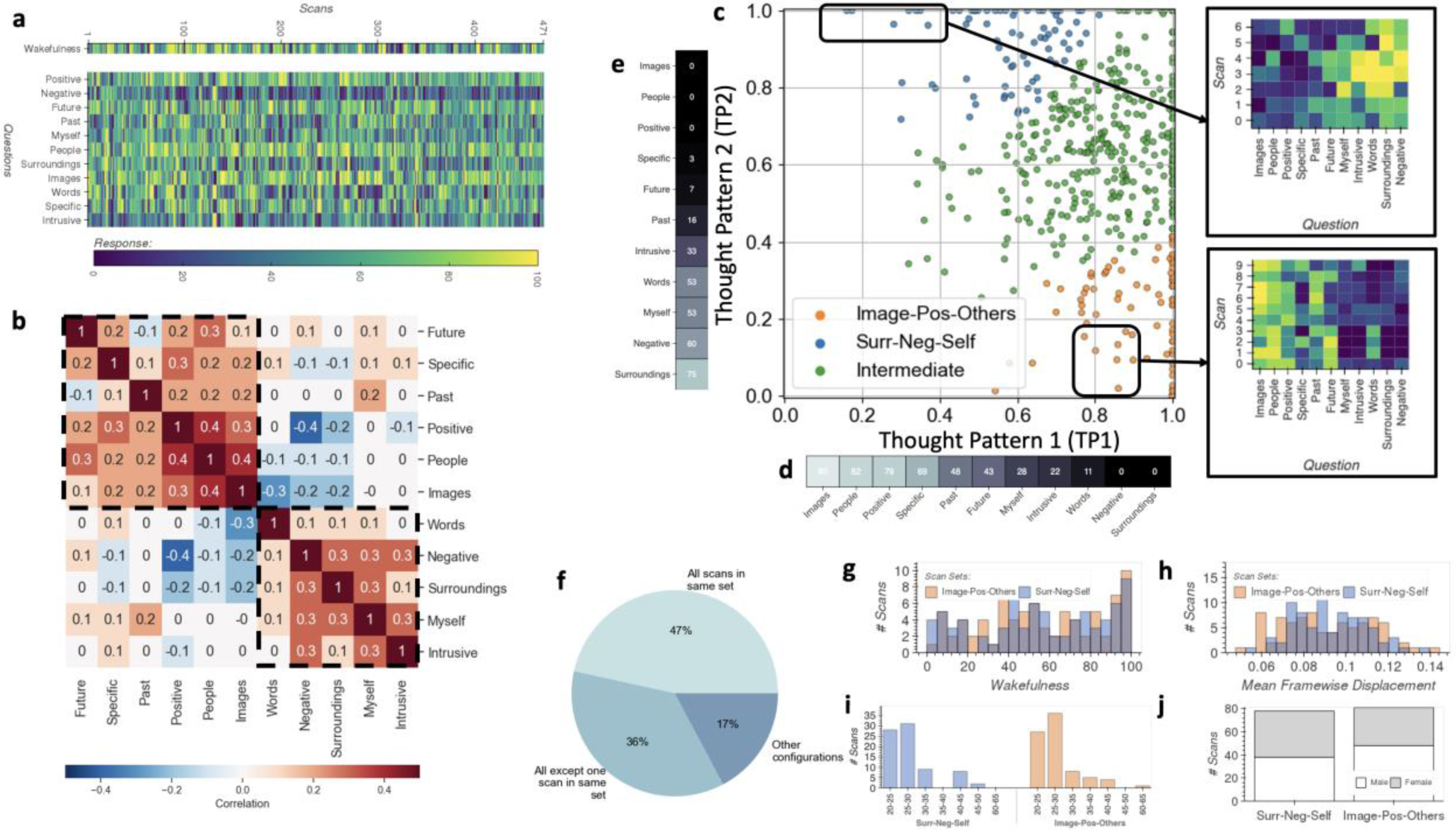
Experiential Data. **a,** Responses to the experiential questionnaire (rows) for all 471 scans (columns). Top row corresponds to the “*wakefulness*” item. The matrix below shows responses to the remaining 11 questions about the form and content of in-scanner thoughts. **b,** Correlation between responses to all questions, except wakefulness. Sets of questions (dashed black outline) appear to correlate more strongly among themselves. **c,** 2D representation of the self-report data (except wakefulness) generated with ICQF^34^, a form of factor analysis. Here, each scan is represented by a colored dot. Axes represent two overall thought patterns extracted by the algorithm: X-axis (Thought Pattern 1 or TP1) represents positive thoughts in the form of images about other people, and Y-axis (Thought Pattern 2 or TP2) represents negative thoughts about the self and the surroundings. Dots representing scans are colored by set membership (agglomerative clustering; number of clusters = 3). To the right of the scatter plot, we depict answers for two groups of representative scans. On the top, we present answers for 7 scans with high loadings for TP2 and low loadings for TP1. High responses concentrate on the questions that more heavily relate to TP2 (e.g., negative, surroundings, words and intrusive). Below that, we show responses to 7 scans with high loadings for TP1 and low loadings for TP2. Here, higher scores concentrate on the images, people, positive and specific items of the questionnaire. **d,** Relationship between TP1 and questionnaire items as a vector of weights outputted by the ICQF algorithm which indicate how strongly each questionnaire item relates to this Thought Pattern axis (top) and as a word cloud (bottom). **e,** Same as d, but for TP2. **f,** Distribution of scans across sets (“Image-Pos-Others”, “Intermediate” or ”Surr-Neg-Self”) for scans acquired on participants that completed a minimum of three resting-state scans. **g,** Distribution of responses to the wakefulness item for the two sets of scans sitting on opposite corners of the 2D space (i.e., *“Image-Pos-Others”* and *“Surr-Neg-Self”*) that will enter the FC analyses based on Network-based Statistics (see Fig. 2). **h,** Distribution of head motion estimates for the same two sets**. i,** Distribution for age range in both sets. **j,** Gender counts in both sets.

Fig. 1b shows the correlation between responses to items about form and content of thoughts, which suggests the presence of two latent axes of variability (dashed rectangles). Based on this observation, we submitted these responses to the Interpretability Constrained Questionnaire Factorization (ICQF) algorithm^34^, which generated a lower dimensional representation of the data in which each scan is now represented as a point in 2D space (Fig. 1c). Because ICQF links original questionnaire items to latent dimensions using only positive weights, one can think of the axes in Fig. 1c as denoting two overall forms of thought (i.e., “*Thought Patterns*”) present in the data. Moreover, because loadings on these axes are constrained to values between 0 and 1, the location of a scan in this 2D space represents how much each of these two Thought Patterns is associated with a given scan. Larger weights on the X-axis (Thought Pattern 1 or TP1) indicate scans strongly associated with thoughts in the form of images, about other people and with a positive valence (Fig. 1d). Larger weights on the Y-axis (Thought Pattern 2 or TP2) indicate scans associated with thoughts focused on the surroundings, self and of negative valence (Fig. 1e).

Several observations are possible using this 2D representation (Fig. 1c). First, the bottom-left corner of the figure is empty despite the adaption of sparsity constraints in the representation, suggesting that questionnaire items surveyed relevant dimensions of in-scanner thought. In other words, thoughts represented, to some degree, by these two overall thought patterns are present in all scans. Second, some scans have high loadings on both dimensions. This is expected given the retrospective nature of the survey: a short questionnaire is used to describe thoughts spanning a 15-minute period during which more than one thought pattern can occur. Third, scans occupy a continuum within the space described by these two axes.

Next, we applied Agglomerative Clustering (*k=3*) to this 2D representation of questionnaire items generated with ICQF as to subdivide the sample into three sets of scans: two sitting on opposite corners of the space (blue and orange in Fig. 1c), and an additional one in-between these two (green). We labeled the first set in the bottom right corner (orange; 81 scans) “*Image-Pos-Others*” as it contains predominantly scans described as consisting of positive thoughts, in the form of images and about other people (i.e., high on TP1 and low on TP2). Next, we labeled the set on the top left corner (blue; 78 scans) *“Surr-Neg-Self”* because it corresponds to scans described as including thoughts focused on the environment, of negative valence or about oneself. The third set, labeled “Intermediate” includes scans sitting in-between the two other groups in the space described by TP1 and TP2. Defining these three sets is a first step towards answering the first two questions targeted in this work: 1) do participants consistently revert to similar thought patterns when asked to rest inside the scanner? and 2) do systematic differences in thought pattern lead to significant differences in rsFC?

One way to reformulate the first question in terms of the sets we just defined is: how often do scans from the same subject belong to the same set? If we consider only scans for subjects who completed a minimum of three resting-state scans (444 scans across 116 subjects), we observe that for 47% of those participants, all scans belong to the same set; for 36% all scans except one are in the same set; and only for 17% of participants scans are either evenly distributed across two sets or separated across the three sets (Fig. 1f). This result suggests that participants tend to, more consistently than not, have thoughts of similar characteristics across consecutive resting-state scans.

### Differences in functional connectivity due to systematic differences in thought patterns

To evaluate the second question concerning whether systematic differences in thought patterns translate into significant differences in FC, we focused our attention on the scans in the two “extreme” sets *(Image-Pos-Others* and *Surr-Neg-Self*) as those are most distinct according to reported thought patterns.

FC matrices for all scans in the *Image-Pos-Others* and *Surr-Neg-Self* sets were created using the 400 ROI/7 network Schaefer Atlas^36^, modified to also include 8 subcortical ROIs from the AAL Atlas^37^ (Fig. 2a, Suppl. Fig. 1). Significant differences in rsFC across these two sets were evaluated using Network-Based Statistics^38^ (NBS; *p_FWE_<0.05*). We found 590 connections (0.82% of all possible connections) involving 221 nodes (58%) to be significantly stronger for the *Surr-Neg-Self* set as compared to the *Image-Pos-Others* set (Fig. 2b-d). These connections tend to be lateralized to the right hemisphere (*FC*_*laterality*_*index*_ = −0.17). No significant connections were found for the inverse contrast. Counts of connections at the network level (Fig. 2d) reveal that the larger proportion of significantly stronger connections correspond to those involving nodes of both the Dorsal and Ventral Attention networks and primary unimodal networks (dotted red square in Fig. 2d). Similarly, we also observe a second cluster of significantly different connections involving the Fronto-Parietal Control network, primary unimodal networks and Attention networks (green dotted rectangle in Fig. 2d).

**Figure. 2.**
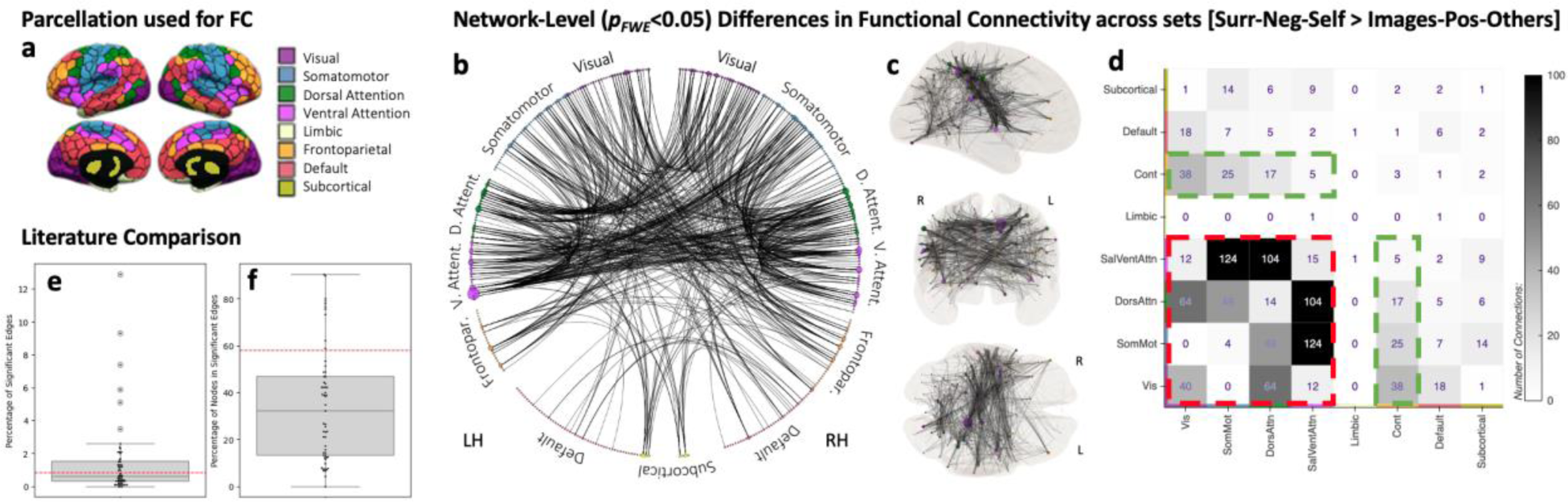
Significant differences in FC across scans with different thought patters. **a,** Schematic of the parcellation used for the generation of FC matrices. We relied on the 400 ROI Shaeffer atlas organized into 7 different networks, as well as a set of additional subcortical ROIs from the AAL atlas. **b,** Circos plot with connections rendered as significant (NBS, 5000 Permutations, Edge-Level Threshold p < 0.001, Network-Level Threshold p < 0.05) for the *Surr-Neg-Self > Images-Pos-Others* contrast. In this plot, ROIs are depicted as colored dots (color indicating network membership) on a circumference, and significant connections are depicted as black lines. ROIs are grouped by network and hemisphere, with left hemisphere ROIs on the left, and right hemisphere ROIs on the right of the plot. The size of the dot is proportional to the degree of the ROI (e.g., number of significant connections emerging from the ROI. **c,** Same information as in b, but this time on a glass brain. ROI size denotes degree and ROI color denotes network membership. **d,** Counts of between- and within-network connections significant for the contrast *Surr-Neg-Self > Images-Pos-Others*. **e,** Boxplot depicting the distribution for percentage of significant edges reported in the NBS literature (see Methods for details on how manuscripts entering these analyses were selected). Colored boxes show quartiles, whiskers extent to show the rest of the distribution, except outliers shown as circles. Individual values contributing to the boxplot are depicted as small black dots. The red dashed line indicates the percentage of connections rendered significant in our study. **f**, Percentage of nodes contributing to significant edges in the selected literature. Boxplot shows the distribution across examined studies, blue dots represent the values for individual contrast and the dotted red line the number of nodes contributing to significant edges in our study.

Because both wakefulness^6,7,33^ and head motion^35^ are known to strongly modulate rsFC, prior to any FC analyses, we checked for differences in these two confounds across the two sets. Fig. 1g shows the distribution of responses to the wakefulness question across the two sets. No significant difference was observed (*T-test=0.16, p=0.87; Mann Whitney U Test=3184, p =0.93*). Fig. 1h shows the same information for mean head motion estimates. No significant difference was observed (*T-test=-0.60, p=0.54; Mann Whitney U Test=2967, p=0.51*). Additionally, we also checked for differences in demographics. Fig. 1i shows scan counts per age range. No significant difference was observed (*Paired T-test=-1.81, p=0.14; Wilcoxon Test=12, p=0.94*). Finally, Fig. 1j shows scan counts by gender. Although not exactly matched across sets, both sets contain similar distributions of male and female subjects.

To evaluate how our results compare to prior clinical work that relies on NBS to uncover clinically relevant rsFC patterns, we calculated the percentage of significant edges (or connections) and nodes (or ROIs) contributing to those in a set of 50 recent scientific papers (see Methods section for details on how these were selected and Suppl. Table 1 for the list of manuscripts). Fig. 2e shows the distribution of percentage of significant edges in the selected manuscripts as a box plot. Our results (0.82%) are marked as a dashed red line. Fig. 2f shows equivalent information in terms of percentage of nodes contributing to significantly different connections. Both figures demonstrate that systematic differences observed in thought patterns can induce significant differences in FC that have spatial extent similar to that previously reported when comparing healthy and clinical populations.

### Prediction of self-reports based on Functional Connectivity

We used the Connectome-based Predictive Modeling (CPM) framework^39^ to evaluate the third question concerning whether it is possible to predict different aspects of in-scanner experience based on rsFC. Using data from all 471 scans, we attempted to predict wakefulness levels, factor scores on TP1 and TP2, and individual responses to the 11 questionnaire items about the form and content of in-scanner thoughts. Prediction accuracy was evaluated in terms of the Pearson’s correlation between observed and predicted values after motion orthogonalization. Significance of the correlation scores were evaluated non-parametrically against null distributions computed on 10,000 randomizations (see Methods for additional details). We were able to significantly predict reported levels of wakefulness, factor scores on both TP1 and TP2, and responses to the Images, Surrounding and Past questionnaire items (*p_non-parametric, uncorrected_ <0.05,* Fig. 3a-c, Table 2). Of these, "Past” did not survive multiple comparison correction based on the False Discovery Rate (FDR^40^) method (*p_non-parametric,_*_FDR_<0.05).

**Figure 3.**
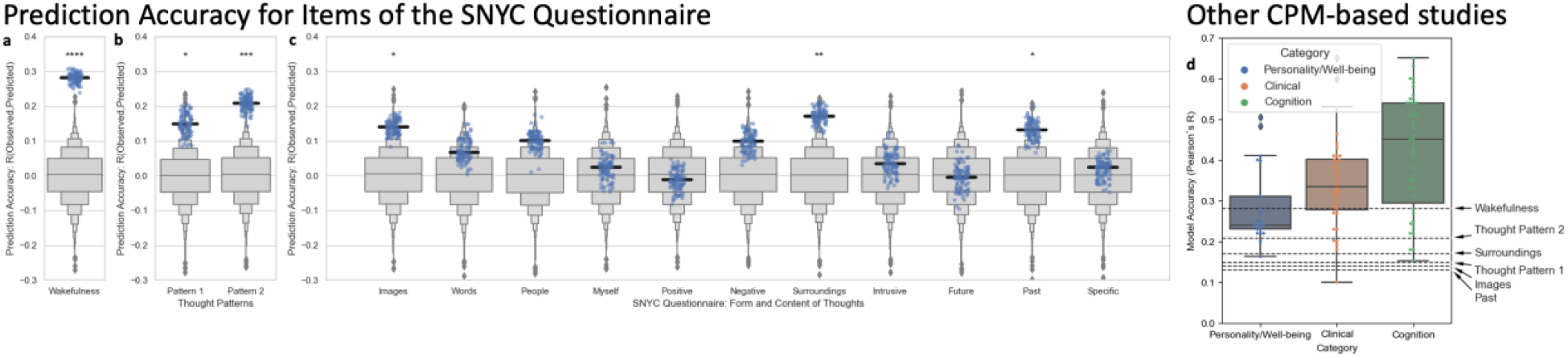
Prediction Accuracy for CPM analyses. **a,** Answers to item *“I was completely awake”*. **b,** *Thought Pattern 1* and *Thought Pattern 2* as defined using the ICQF algorithm. **c,** Answers to the 11 items about the form and content of in-scanner thoughts. In all plots, the gray boxenplot depicts the null distribution of accuracy values across 10,000 randomized permutations. Blue dots correspond to the Pearson’s correlation between observed and predicted values (i.e., prediction accuracy) in each of the 100 iterations used to predict each target. Bold black lines represent the median prediction accuracy across the 100 iterations. Significance annotations correspond to the non-parametric, uncorrected p-value of the median prediction accuracy according to each null distribution (*\*\*\*\* p<0.0001, *** p<0.001, ** p<0.01, * p< 0.05*). **d,** Comparison to previously published studies that used CPM to predict cognitive, personality or clinical traits based on full-brain functional connectivity. For each category of studies, we provide a boxplot depicting the distribution of reported accuracy values (colored boxes show quartiles, whiskers extent to show the rest of the distribution, except outliers shown as diamonds). Colored dots represent all individual accuracy values contributing to each distribution. In addition, horizontal dashed black lines show accuracy for the 6 models rendered significant in this study. Details on how we conducted the literature search contributing to this panel can be found on the online methods. Suppl. Table 2 list the studies contributing to this figure panel.

**Table 2.**
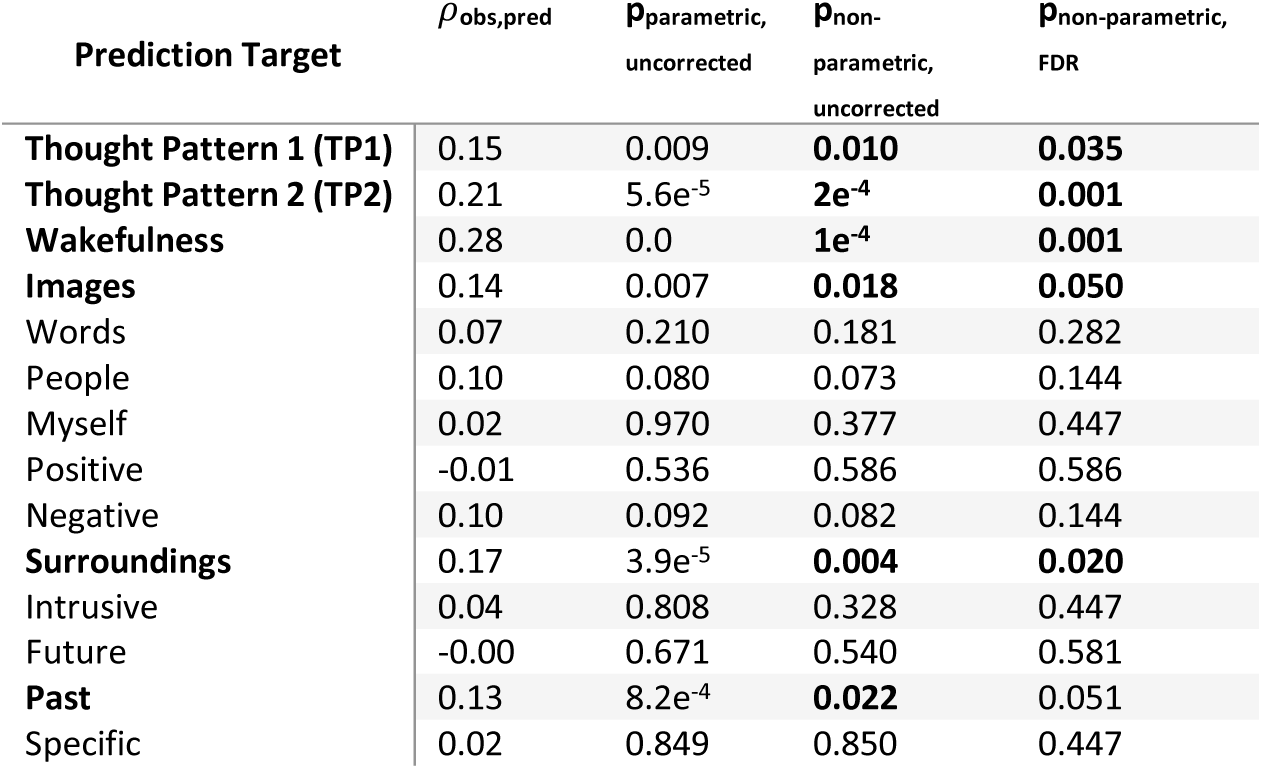
Model accuracies for all 14 CPM models. *ρ_obs,pred_* = Median Pearson’s correlation between observed and predicted values across all 100 permutations. *p*_parametric,uncorrected_ = parametric and uncorrected p-value. *p*_non-parametric,uncorrected_ = non parametric, uncorrected p-value based on the 10,000 null permutations generated separately for each prediction target. *p*_non-parametric,FDR_ = non parametric, FDR corrected p-value. p-values < 0.05 highlighted in *Bold* font.

| Prediction Target | $\rho_{obs, pred}$ | $p_{parametric, uncorrected}$ | $p_{non-parametric, uncorrected}$ | $p_{non-parametric, FDR}$ |
| --- | --- | --- | --- | --- |
| **Thought Pattern 1 (TP1)** | 0.15 | 0.009 | **0.010** | **0.035** |
| **Thought Pattern 2 (TP2)** | 0.21 | 5.6e<sup>-5</sup> | **2e<sup>-4</sup>** | **0.001** |
| **Wakefulness** | 0.28 | 0.0 | **1e<sup>-4</sup>** | **0.001** |
| **Images** | 0.14 | 0.007 | **0.018** | **0.050** |
| Words | 0.07 | 0.210 | 0.181 | 0.282 |
| People | 0.10 | 0.080 | 0.073 | 0.144 |
| Myself | 0.02 | 0.970 | 0.377 | 0.447 |
| Positive | -0.01 | 0.536 | 0.586 | 0.586 |
| Negative | 0.10 | 0.092 | 0.082 | 0.144 |
| **Surroundings** | 0.17 | 3.9e<sup>-5</sup> | **0.004** | **0.020** |
| Intrusive | 0.04 | 0.808 | 0.328 | 0.447 |
| Future | -0.00 | 0.671 | 0.540 | 0.581 |
| **Past** | 0.13 | 8.2e<sup>-4</sup> | **0.022** | 0.051 |
| Specific | 0.02 | 0.849 | 0.850 | 0.447 |

To predict Wakefulness, CPM relied on a model that includes 3839 connections (5.3% of all possible connections) distributed across all seven networks (Fig. 4a): 2026 (2.8%) of these connections exhibit positive correlation with the prediction target, and the remaining 1813 (2.5%) show negative correlation. A large fraction of the positively correlated connections represent links between subcortical and cortical regions (Fig. 4b-c). Conversely, connections that are negatively correlated with Wakefulness are cortex-to-cortex connections involving primarily the Default, Ventral and Dorsal Attention, and primary unimodal networks (Fig. 4d-e).

**Figure 4.**
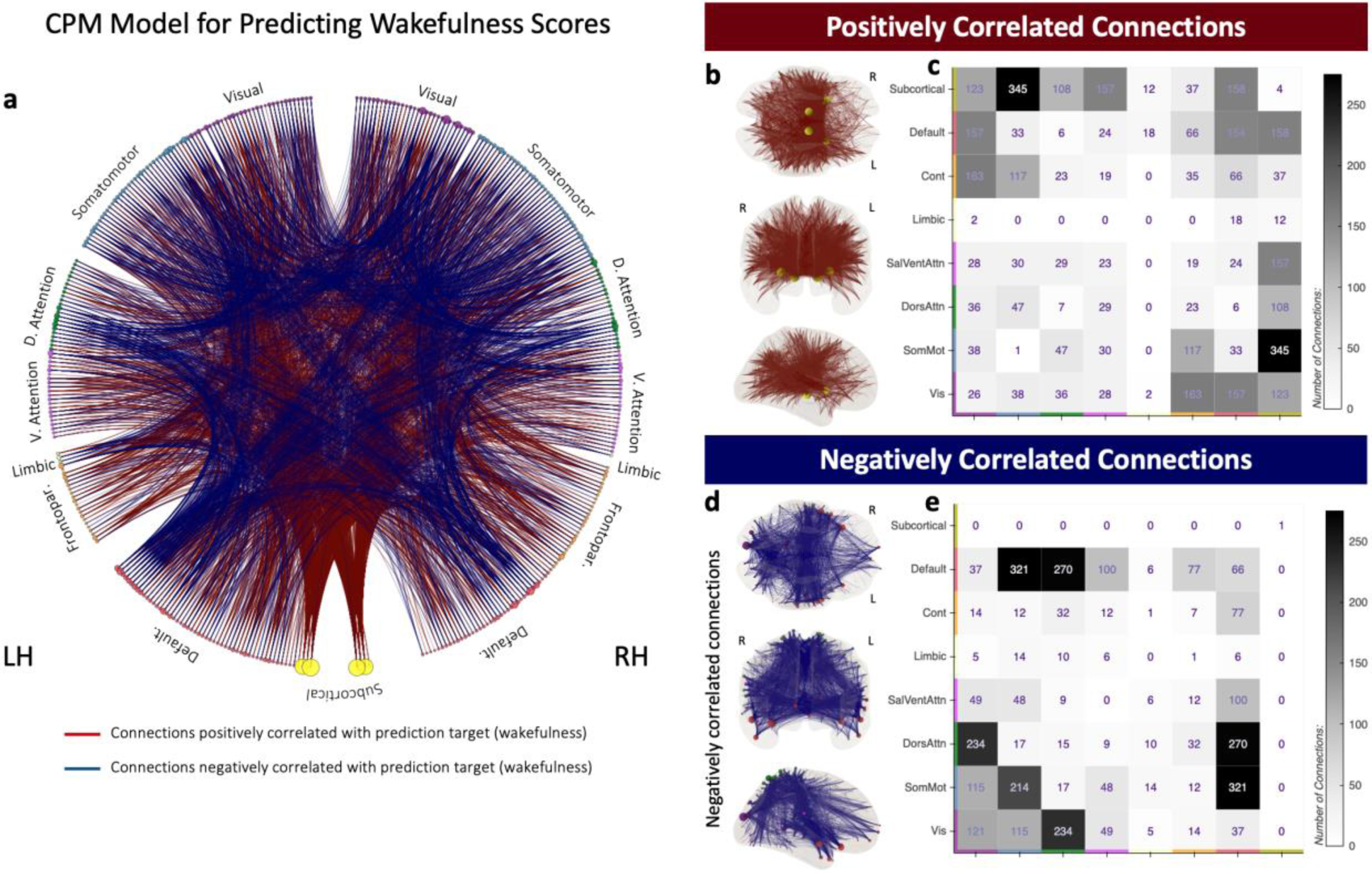
CPM model for Wakefulness item *“I was completely awake”*. **a,** Circos plot depicting connections found to correlate positively (dark red) and negatively (dark blue) with the prediction target. **b,** Connections positively correlated with reported levels of Wakefulness on a glass brain. **c,** Counts of within- and across-network connections that correlate positively with reported levels of Wakefulness. **d,** Connections negatively correlated with reported levels of Wakefulness on a glass brain. **e,** Counts of within- and across-network connections that correlate negatively with reported levels of Wakefulness. In **a,b,** and **c,** the size of the circles representing regions of interest (ROI) is proportional to the number of connections that include the ROI. The color of the circles depicts network membership.

Next, Fig. 5 shows the CPM models for predicting factor scores on the two Thought Patterns previously described. The CPM model for TP1 (Fig. 5a-e) is sparser (1036 connections | 1.4%) than the one for TP2 (1778 connections | 2.5%; Fig. 5f-j). For TP1, positively correlated connections involve primarily nodes of the Frontoparietal Control and Default networks (black dashed rectangles in Fig. 5b); with highest degree nodes sitting in bilateral dorsolateral prefrontal cortex (black arrows in Fig. 5c). Conversely, connections that correlate negatively with TP1 tend to concentrate in central and posterior areas of the brain that belong primarily to the Visual and Dorsal Attention networks. Highest degree nodes for this second portion (negative correlations) of the model include the left medial intraparietal sulcus, part of the Dorsal Attention network, and left para-hippocampal area, part of the Visual network (black arrows in Fig. 5e).

**Figure 5.**
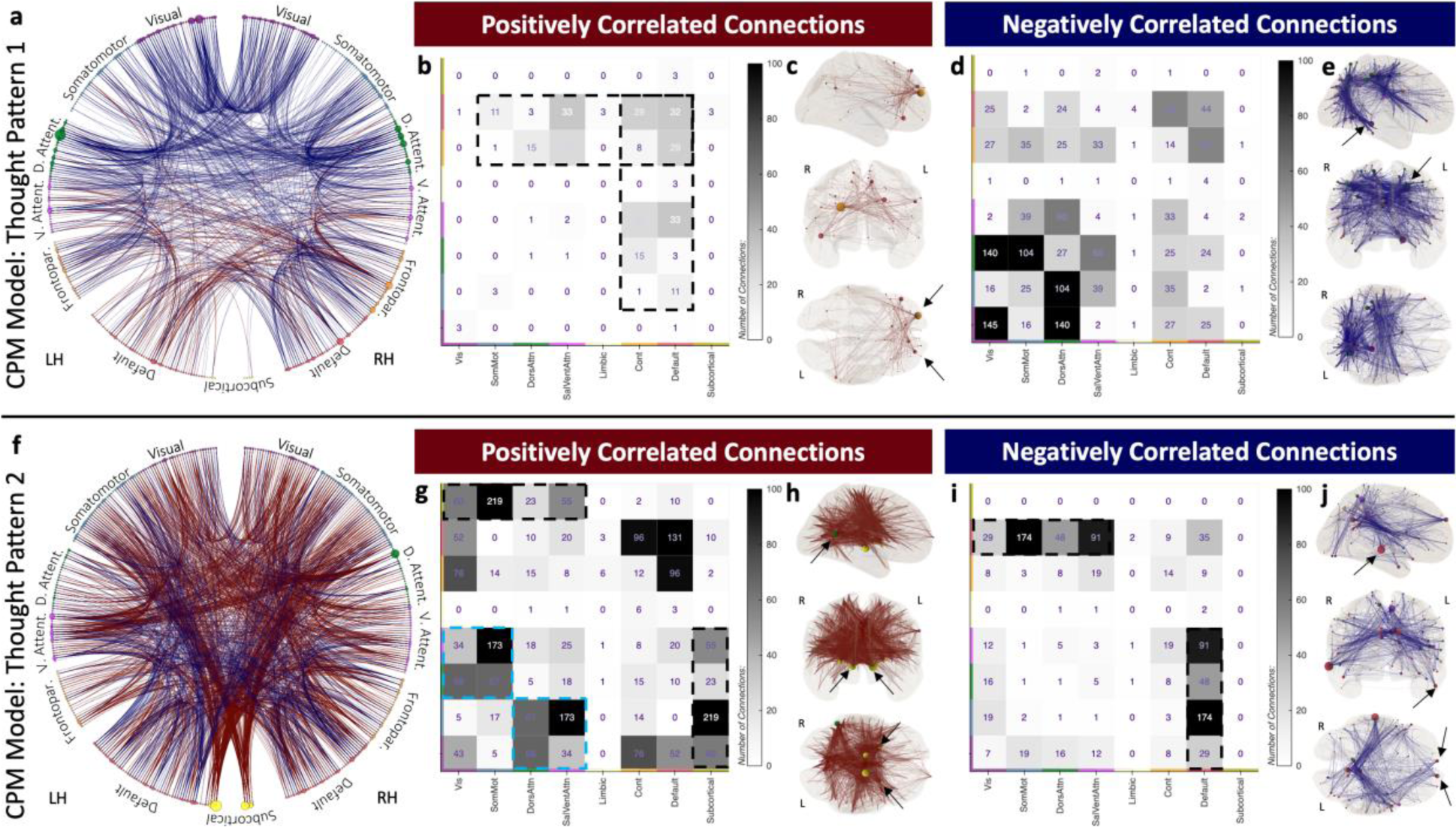
Prediction Model for Thought Patterns 1 and 2. **a,** Circos plot of the CPM model for the prediction of loadings on TP1 axis. In all panels dark red lines and dark blue lines denote, respectively, connections that correlate positively or negatively with the prediction target. **b,** Counts of within- and across-network connections that correlate positively with TP1. **c,** Connections positively correlated with TP1 on a glass brain. **d,** Counts of within- and across-network connections that correlate negatively with TP1. **e,** Connections negatively correlated with TP1 on a glass brain. **f-j,** Equivalent results to **a-e** but using as prediction target TP2 loadings instead of TP1. The color of circles representing ROIs in Circos plots and glass brains denote network membership, and the size of the circle the number of connections that include a given ROI (e.g., its degree).

For TP2, the CPM model has 1269 (1.8%) positively correlated connections widely distributed across all networks, except the Limbic network. That said, larger connection counts occur for connections between subcortical regions and regions from both attentional networks and primary unimodal networks (black dashed rectangle in Fig. 5g), as well as between both attentional networks and primary unimodal networks (cyan dashed rectangle in Fig. 5g). Similarly, top degree nodes include bilateral thalami and putamen, as well as the right temporo-parieto-occipital junction, which is part of the Dorsal Attention network (black arrows in Fig. 5h). Regarding the 509 (0.7%) connections that negatively correlate with TP2, these are primarily connections between nodes of the Default network and nodes in both Attention and unimodal primary networks (black dashed rectangle in Fig. 5i). In this case, highest degree nodes are located primarily in the Default network (black arrows in Fig. 5j).

Finally, Figure 6 shows the CPM models for the three individual questionnaire items about the form and content of thoughts that the model significantly predicted, namely: Images, Surroundings and Past. The CPM model for Images (Fig. 6a-b) shares features with that of TP1 (Fig. 5c,d), of which Images is the most prominent characteristic (Fig. 1e). Common features between the Images and TP1 models include a tendency for positively correlated connections (dark red connections, Figs. 5c and 6a) to involve regions in dorsolateral prefrontal cortex and negatively correlated connections to concentrate on posterior aspects of the brain, primarily in occipital cortex (dark blue connections, Figs. 5.e and 6b). Some level of agreement was also observed for the CPM model for Surroundings (Fig. 6c-d) and that of TP2 (Fig. 5h,j), of which Surroundings is the most prominent characteristic (Fig. 1d). In this instance, similarities across models include the heavy contribution of subcortical regions to positively correlated connections, and of Default network nodes to their negatively correlated counterparts. For Past, which was not as strongly associated with either Thought Pattern, we observe a unique CPM model primarily composed of positively correlated edges that has its two higher degree nodes located in left frontal regions of the Default network (Fig. 6e).

**Figure 6.**
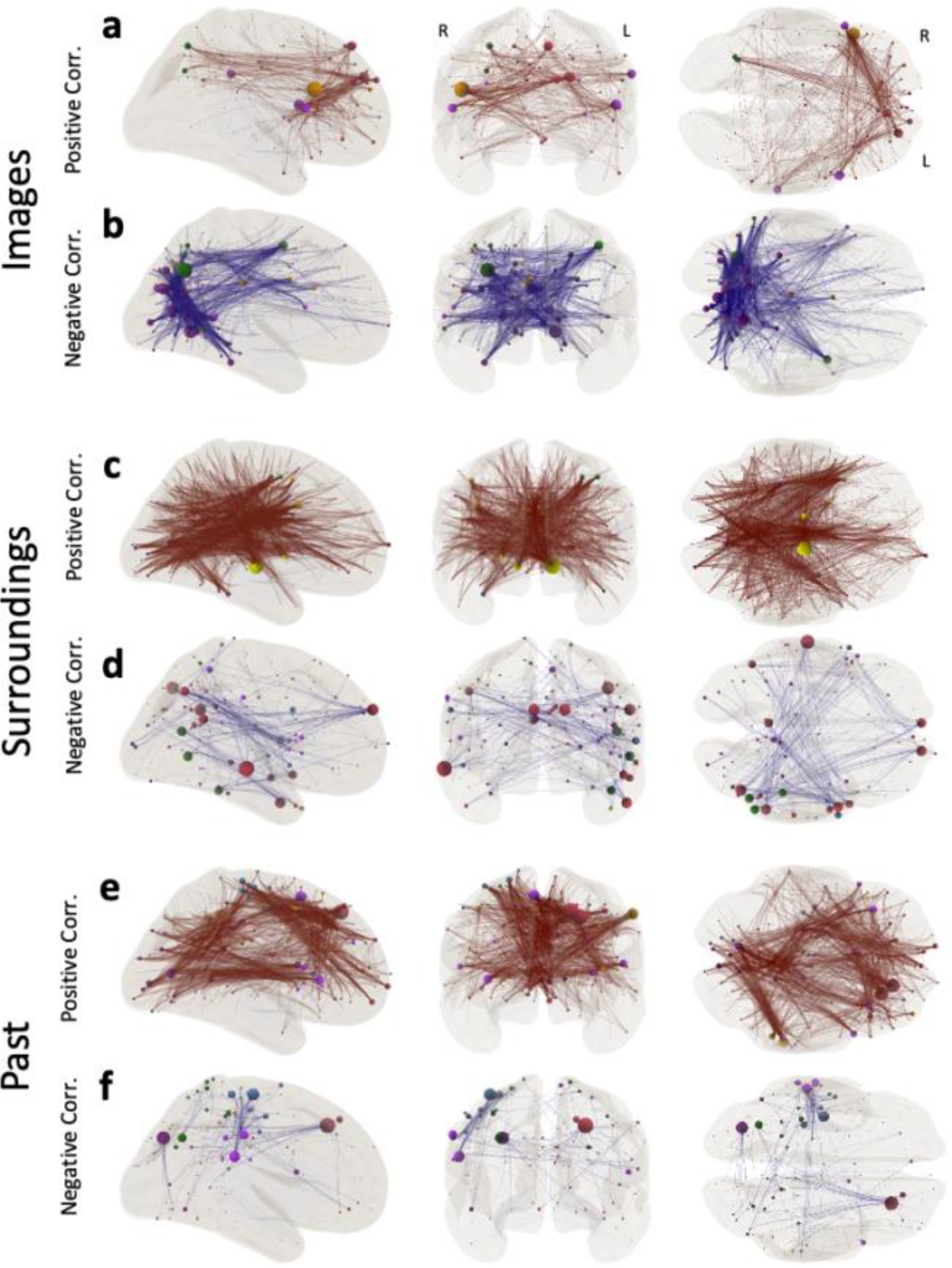
CPM models for individual items related to the form and content of thought that we could predict above significance. **a-b,** Predictive model for Images. **c-d,** Predictive model for Surroundings. **e-f,** Predictive model for Past. In all instances, positively correlated connections are shown on the top row (**a,c,e**) in dark red, and negatively correlated connections on the bottom row (**b,d,f**) in dark blue. Node size denotes degree and node color denotes network membership (see Fig. 2a)

## Discussion

This investigation provides strong evidence for the key—yet often overlooked—role that subjective in-scanner experience plays in shaping resting-state FC. Our results demonstrate the enhanced research value of annotating rs-fMRI data with experiential information to facilitate a more direct exploration of what aspects of intrinsic activity during rest are related to specific attributes of self-generated thoughts. Importantly, this study offers three unique contributions: 1) subjects have similar experiences across successive rs-fMRI sessions, 2) systematic differences in in-scanner experiences yield inter-population differences in resting-state FC, 3) resting-state FC can predict some aspects of in-scanner experiences. These findings are relevant to those studying the neural correlates of spontaneous thought, and more generally, to those that use rs-fMRI to map the functional connectome of healthy and diseased populations. We discuss each of these contributions and their implications below.

First, we show that for subjects with multiple rsfMRI scans in our sample, those scans are often grouped together when we divide the sample according to how scans load on TP1 and TP2 (Figure 1f). This suggests that subjects fall back on similar thought patterns—at least at the level of detail captured by the sNYCQ—every time they participate in a resting-state session. This agrees with prior literature on the topic. For example, Diaz et al.^41^, using the Amsterdam Resting-State Questionnaire (ARSQ), found a high level of test-retest reproducibility for the seven dimensions of experience captured by that tool (Discontinuity of Mind, Theory of Mind, Self, Planning, Somatic Awareness, Comfort, and Sleepiness) across two resting-state scans that occurred at the beginning and end of a multi-task scanning session. In a follow-up study^42^, using an updated version of the ARSQ that also quantifies Health Concerns, Visual Thought and Verbal Thought, the authors found stable and significant test-retest reliability for intervals of 3–32 months in all explored dimensions of in-scanner experience except for Sleepiness and Health Concerns. These reports, combined with ours, suggest that subjects default back to overall similar thought patterns and feelings inside the scanner, suggesting that summary characteristics of in-scanner experience captured with retrospective tools should be considered trait-like, as opposed to state-like, characteristics given their long-term temporal stability.

This finding has important interpretational and analytical implications. First, the high within-subject test-retest reliability of rsFC has often been used as an argument against the possibility of “resting cognition” being a contributing factor to rsFC under the assumption that “resting cognition” would vary from scan to scan^3,43^. While our results, as well as those of others discussed in the previous paragraph, do not imply that subjects have the same exact thoughts and feelings during different scan sessions, it demonstrates that overall high-level descriptors of in-scanner experience (such as those captured by sNYCQ items) might be more static and reproducible within-subject than originally assumed. In fact, this propensity of subjects to have similar thought patterns every time they enter the MRI scanner can explain why in-scanner experience being a modulatory factor of rsFC is compatible with rsFC having high test-retest within-subject reliability. Second, these results also imply that simple within-subject, across-scan averaging will not be efficient at removing in-scanner experiential effects, should that be desirable. A more effective alternative might be to gather experiential information using tools such as the sNYCQ or the ARSQ^41,42^, so that it can be included as nuisance factors in analytical models (similar to how motion and other physiological measures are often incorporated in rsfMRI studies).

Our second research question aimed to quantify whether in-scanner experience is associated with resting-state FC patterns, thereby posing a confound (or an additional explanatory factor) for studies that compare rsFC across populations–a ubiquitous approach when looking to uncover clinically relevant FC patterns. Our analyses show significant differences in rsFC between data from two sets of healthy controls (matched for head motion, age, gender, and wakefulness levels) that differ in terms of the content and form of their in-scanner spontaneous thoughts (Figure 3b-d). Importantly, the spatial extent of such differences—as quantified by percentage of regions and connections—is equivalent to that of clinical studies based on the same analytical methods (Figure 3.e-f). This observation suggests that systematic inter-population differences in what subjects feel and think while being scanned can affect interpretation in population-contrasting studies. Yet, the severity and interpretational challenges will be study-specific as they depend on (1), what aspects of spontaneous thoughts/experience are different across contrasted populations (e.g., overall valence, form, attentional focus, etc.) and (2), the spatial overlap between the connections that are modulated by the study inclusion criteria (e.g., presence/absence of a given clinical condition) and those due to in-scanner experience.

Beyond wakefulness (which has been extensively discussed elsewhere^6,7,33^), one aspect of in-scanner experience that we also found to be key is whether subjects focus their attention on their surroundings. Significantly stronger connections for scans with high loadings on TP2—which is most strongly associated with the Surroundings item of the sNYCQ (Figure 1e)—involve primarily nodes of the Salience/Ventral Attention Network, the Dorsal Attention Network and primary somatosensory networks (red dashed rectangle in Figure 2d). This pattern of stronger FC among these networks is compatible with the well-established role that these networks and their interactions play in the reorientation of attention towards stimuli^44,45^. Similarly, the observed right hemisphere lateralization of connections that survived the *“Surr-Neg-Self > Images-Pos-Others”* contrast is consistent with the known right dominance of spatial attentional processes^46,47^. Such compatibility between our observations and well-established functional roles for the regions and networks involved further supports the claim that the differences in rsFC observed here are indeed due to differences in in-scanner experience across scan sets.

Additionally, our findings provide novel insights regarding the neural correlates of spontaneous thought. For example, the highest degree region for the *“Surr-Neg-Self > Images-Pos-Others”* contrast was region *7Networks_LH_SalVentAttn_Med_5* from the 7Networks/400ROI version of the Schaefer atlas^36^. This region sits on the posterior end of the cingulate sulcus, overlapping primarily with Broadman Area 23c, which is commonly activated by tasks that require an external focus^48^. Prior studies have demonstrated that Area 23c is also involved in valence modulation of nonintentional spontaneous thoughts^49,50^. Using music to experimentally modulate the valence of spontaneous thoughts, these prior studies found Area 23c to increase its centrality during negative thoughts^49,50^. Our results, which highlight the involvement of this region in scans with high loadings on TP2—which is also strongly associated with negative thoughts (Figure 1e)— confirm these prior studies. They also extend them by showing that the involvement of this region in negative spontaneous thoughts is independent of the presence of any modulating stimuli (i.e., the mood inducing music).

Finally, we show that it is possible to predict aspects of in-scanner experience—namely Wakefulness, Images, Surroundings, Past, TP1 and TP2—using rsFC matrices. This further supports the idea that what subjects feel and think during rsfMRI scans leaves a reliable and detectable imprint on rsFC. When we compare our accuracy estimates to studies that use CPM to predict clinical, cognitive and personality phenotypes (Suppl. Tables 2 and 3) we observe that our six models—despite satisfying individual criteria for statistical significance, and five of them surviving FDR-based correction—fall along the lower end of the distributions of accuracies reported in the literature (Fig. 3d). This can be interpreted in at least two ways. It could be argued that this observation suggests that in-scanner experience has a weaker signature in rsFC than phenotypes such as the severity of clinical symptoms, a person’s cognitive abilities or personality traits. Alternatively (or additionally), lower accuracy could be the consequence of the imprecision and limited scope of retrospective introspection instruments for capturing relevant aspects of in-scanner experience during minutes-long rsfMRI scans (see Gonzalez-Castillo et al.^2^ for a detailed discussion). Discerning between these two interpretations requires additional rsfMRI data with more accurate and extensive experiential annotations. Those could be obtained by combining first-person reports with in-scanner recordings (e.g., pupil size, gaze, skin conductance, videography^51^) that also carry information about subjects’ state of mind (e.g., stress and vigilance levels, focus of attention, occurrence of basic emotions, etc.). With such additional data, researchers will be able to evaluate whether accuracy improves as higher quality experiential information becomes available, and perform external across-dataset validation of CPM models, which is the optimal way to evaluate the generality of these models whenever possible^52^.

That said, and as we observed in the contrast analysis (Question 2), topological properties of our CPM models are reconcilable with current models of brain function. For example, the model for Wakefulness (Figure 4a) being the denser of all models is compatible with the large functional reconfiguration that accompanies descent into sleep^6^. Thus, this CPM model includes many subcortical-to-cortical connections that positively correlate with Wakefulness (Figure 4b-c) and many cortical-to-cortical connections that correlate negatively with Wakefulness (Figure 4d-e). This pattern agrees with sleep onset being associated with reduced subcortical-to-cortical connectivity and increased cortical-to-cortical connectivity^6,53^. Also, the CPM model for TP2 recapitulates (Figure 5g) the involvement of both attention networks and primary somatosensory networks observed in the contrast analysis (Figure 2b-d). Finally, one prominent characteristic of the Images and TP1 models is the many within-visual connections that correlate negatively with reported levels of visual imagery (Figures 6-ab & 5a-e). Prior work has shown that within visual network connectivity decreases when subjects engage in continuous visuospatial attention tasks^54,55^ and during natural movie viewing^56^. While the role of visual areas in visual imagery remains controversial (traditional views advocate in favor^57^, but recent meta-analytic^58^ and lesion studies suggest otherwise^59^), our results suggest that extended periods of visual imagery have a similar segregating effect within the visual network during rest to that of attention towards visual stimuli during active tasks.

In summary, this work shows that what subjects think and feel during rsfMRI scans matters. It extends prior work showing how seemingly irrelevant differences in pre-scanning instructions (e.g., ‘let your mind wander’ vs. ‘think of nothing’^60^) can alter FC patterns as well as past studies demonstrating that in-scanner experience and spontaneous thought patterns can be related to resting-state signals^11,12,61^. It highlights how introspective reports can enhance the interpretation of rsfMRI by attaching a meaningful neurocognitive interpretation to findings of differing rsFC. Importantly, we do not claim that in-scanner experience and spontaneous thought patterns are the only (or primary) factor driving rsFC. As briefly mentioned in the introduction, other neurobiological and cognitive processes (e.g., memory consolidation, homeostatic processes, etc.) might be equally (or more) important. Future research should elucidate the relative contributions of these and other factors. Similarly, our results should not be considered grounds for diminishing rsfMRI in favor of alternative experimental designs such as naturalistic protocols. While naturalistic stimuli can induce synchronized neural responses across subjects (offering analytical approaches not available to rsfMRI^62^), subjects might still experience the same movie (or audio story) differently on repeated presentations (e.g., by focusing on different aspects, recalling prior presentations, or eliciting different emotions), and therefore accurate interpretation of naturalistic-based results might also ultimately require experiential annotations, as we propose for resting-state.

Resting-state fMRI occupies a central place in neuroimaging research, yet its inability to fulfill clinical standards (i.e., sufficient specificity and sensitivity) and often unsatisfactory interpretation have stalled progress in recent years. Here, we show that in considering the role that in-scanner experience plays in shaping rsFC might address these issues by accounting for unexplained variance in the data. This is not an insurmountable challenge. Much of that variance might be properly modeled once we learn what aspects of experience matter most, their co-occurrence with different clinical conditions and how to accurately measure their presence/intensity during fMRI scans. Ultimately, this work suggests that is possible to move beyond the current impasses if we, as a community, are willing to embrace and help resolve the many complexities that arise when incorporating first-person data into our research agendas.

### Online Methods

#### Dataset

This work is conducted using portions of the publicly available Max Planck Institute Leipzig Mind-Brain-Body dataset^27^. This dataset includes, among other items, resting-state scans (15.5 mins long, 2.3×2.3×2.3mm, TR = 1.4s, 3T) annotated with retrospective self-reports of information regarding the content and form of thoughts participants had during the scan (see Table 1). For each resting-state scan, it also contains information about participant’s subjective perception of how awake they remained during the scan, as well as age range (in 5-year intervals) and gender. All information about scanning parameters, demographics, and behavioral tests included in this dataset can be found in the Scientific Data publication that accompanies the release of the dataset^27^.

The original dataset contains a total of 788 resting-state scans acquired on 194 participants. The number of scans per participant varies between one and four. Scans are distributed across a maximum of two different scanning occasions. Only 693 scanned (distributed across 175 participants) had complete annotations about the form and contents of thoughts, as well as subjective reports of wakefulness. It is those 693 that we selected for our analyses.

#### fMRI Data Pre-processing

For the 175 participants that had at least one resting-state scan annotated with in-scanner experience data, we attempted preprocessing of their T1 scans using the anatomical pre-processing pipeline distributed with the dataset (https://github.com/NeuroanatomyAndConnectivity). This pipeline includes the following operations: removal of T1-weighted image background (CBS Tools), cortical surface reconstruction (Freesurfer), and registration towards the MNI152 1mm template (ANTs). For 25 subjects, the pipeline did not finalize successfully (e.g., errors in tissue segmentation, skull stripping, etc.). Resting-state scans from these subjects (191 scans) were not considered in the rest of this work (Table 3).

**Table 3.** Number of scans discarded at each pre-processing and QC step.

| *Elimination Criteria* | *Rest Scans Removed* | *Scans Remaining* | *Subjects Remaining* |
| --- | --- | --- | --- |
| *Short NYCQ missing* | 95 | 693 | 175 |
| *Anatomical Preprocessing Failed* | 96 | 597 | 150 |
| *Functional Preprocessing Failed* | 13 | 584 | 147 |
| *Excessive Head Motion* | 113 | 471 | 133 |

Pre-processing of the remaining 597 resting-state scans proceeded with two steps. First, we used the initial portion of the functional pre-processing pipelines distributed with the MPI dataset to perform the following operations: discard initial five volumes, head motion correction, spatial distortion correction, co-registration to MNI space. Next, we performed the following three additional steps using AFNI^63^: temporal scaling by dividing by the mean, nuisance regression (motion, first derivative of motion, Legendre polynomials up to third degree, physiological noise regressors computed with the aCompCor method^64^), and temporal filtering [0.01 – 0.15 Hz]. Functional pre-processing failed for 13 resting-state scans due to missing functional or field map data (Table 3).

For scans that completed the functional pre-processing pipeline, we used traces of relative displacement to mark as invalid volumes with a relative displacement above 0.3mm. For any such volume, we also marked as invalid the preceding volume and the following two. Any scan with 30% or more of volumes marked as invalid was discarded from any additional analyses. This resulted in the removal of an additional 113 scans. Table 3 below summarizes the number of scans rejected at every pre-processing step.

#### In-scanner Experience Reports Analyses

##### Overview

In-scanner experience data was divided in two parts: one contained responses to the 11 questions about the form and content of in-scanner thoughts, and the second was a vector with the response to the wakefulness item (Fig. 1a). The dimensionality reduction and subsequent agglomerative clustering steps described below were conducted using only the first part (i.e., answers to all questions except the one about wakefulness). Answers to the wakefulness item were only used to evaluate potential differences in wakefulness levels across scan sets generated via agglomerative clustering and as a target for prediction (see Connectome Predictive Modeling section below).

##### Dimensionality Reduction

Answers regarding the form and content of in-scanner thoughts (matrix *M*_471 scans X 11 questions_; Suppl. Fig. 2a) and demographic information (matrix *C*_471 scans X 5 demographic encoding variables_; Suppl. Fig. 2c) were input to the Interpretability Constrained Questionnaire Factorization (ICQF)^34^ algorithm to generate a low dimensional representation of the data. We chose this matrix decomposition technique, as opposed to Principal Component Analysis, because it is better suited for distributions that are both non-negative and non-Gaussian (Suppl. Fig. 3 shows that was the case for these data), ensure positive loadings in the low dimensional space (which is key for interpretational purposes), and account for potential confounding effects between demographics and questionnaire items.

In a nutshell, the ICQF algorithm attempts the following matrix decomposition:

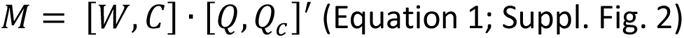

where *M* and *C* are the inputs to the algorithm described above. *W* is the resulting low dimensional representation (*d* = number of dimensions) of the data (471 scans X *d* Dimensions) with entries constrained to the range [0,1]. *Q* is a 11 questions X *d* Dimensions matrix with values constrained to the range [0,100] and provides information about how each dimension in the lower space relates to the original 11 questionnaire items. *Q*_*c*_ is a 11 question X 5 demographic items matrix with values also in the range [0,100] that contains information about how age and gender relate to the original questionnaire items. Finally, as stated above, C is a 471 x 5 matrix with demographic information encoded as follows: the first column models an intercept (all ones), the next two columns encode age range using hashing encoding constrained to the [0,1] range, and the last two columns encode gender using one hot encoding. In Equation 1 above, [*A*, *B*] denotes horizontal concatenation of matrices *A* and *B*.

Given *M* and *C*, the ICQF algorithm finds *Q*, *Q*_*c*_ and *W* using stochastic gradient descent and an Alternating Direction Method of Multipliers during optimization (see Lam et al.^34^ for additional details). The ICQF algorithm has three hyper-parameters, namely the number of dimensions (*d*), the sparsity level for the *W* matrix (*β*_*W*_), and the sparsity level for the *Q* matrix (*β*_*Q*_). We used cross-validation to estimate (in a data-driven manner), the optimal values for these three parameters. The range of values explored for each hyper-parameter and optimal values selected are reported in Table 4. Cost functions associated with these exploratory analyses are shown in Suppl. Fig. 3.

**Table 4.** Hyper-parameter Exploration for the *ICQF* algorithm.

| HYPER-PARAMETER | DESCRIPTION | EXPLORATION RANGE | OPTIMAL VALUE |
| --- | --- | --- | --- |
| $d$ | Number of dimensions | 1,2,3,4 | 2 |
| $\beta_W$ | Sparsity for W matrix | 0.0, 0.01, 0.1, 1, 2, 3 | 0.0 |
| $\beta_Q$ | Sparsity for Q matrix | 0.0, 0.01, 0.1, 1, 2, 3 | 0.01 |

##### Agglomerative Clustering

We grouped scans into three sets using the scikit-learn^65^ implementation of the Agglomerative Clustering Algorithm. Inputs to the algorithm were the matrix *W* and *k* = 3 (i.e., number of desired sets). We used *k* = 3 because our goal was not to uncover well delineated clusters (which were not observed and would have required a data driven estimate of *k*) but simply to subdivide the sample into three sets: two with items sitting on the corners of the low dimensional space, and one containing all other items in the sample.

#### Functional Connectivity Analyses - Statistical Differences across scan sets

##### Parcellation

Following pre-processing, we computed representative time-series (spatial mean across voxels) and FC matrices separately for each scan using a modified version the 400ROI / 7 Networks Schaefer atlas^36^ using the AFNI^63^ program *3dNetCorr*. Thirteen ROIs were removed from the atlas prior to generating the FC matrices because they include regions suffering from excessive signal dropout. Those correspond to the 12 regions forming the Limbic network and one region part of the default mode network labeled as “*LH_Default_Temp_2*”. Additionally, 8 subcortical regions for the AAL atlas^37^ (bilateral thalami, putamen, striatum, and pallidum) were added. Suppl. Fig. 1 shows the final version of the atlas used in the analyses.

##### Network-Based-Statistics: Significant Testing

We used the Network-Based Statistic (NBS^38^) procedure to identify significant differences in FC between scan sets on opposite ends of the in-scanner experience space defined above. The NBS algorithm proceeds as follows. First, the procedure identifies individual edges that surpass an edge-level significance level of p < 0.001 using a two-sample T-test on a design matrix (Suppl. Fig. 5) that encoded both set membership (e.g., *Images-Pos-Others* or *Surr-Neg-Self*) and subject identity, to account for the fact that there were repeated scans per subject in the sample. Next, the algorithm looks for connected components of supra-threshold edges and computes their extent (i.e., size). This is a network equivalent of how cluster size is computed on supra-threshold statistical activity maps. Third, a null distribution of connected component sizes is generated using 5,000 permutations of scan labels. This null distribution is used to evaluate the statistical significance of observed connected sets of supra-threshold edges. Only those with a component-level p < 0.05 are rendered significant and retained for interpretation, which is equivalent to family-wise error corrected significance for the identified network components.

##### Laterality Index

To quantitatively evaluate if connections involve primarily regions from one hemisphere over the other, we computed the FC Laterality Index (*FC*_*laterality*_*index*_) using the following formula:

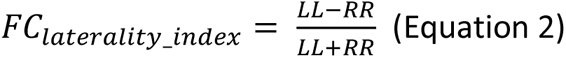

where *LL* is the number of left intra-hemispheric significant connections and *RR* is the number of right intra-hemispheric significant connections. The *FC*_*laterality*_*index*_ ranges between -1 (right-dominant) to 1 (left-dominant).

This formula was adapted from its equivalent standard formulation for activity-based results^66,67^ by substituting active-voxel counts by number of significant connections.

#### Functional Connectivity Analyses – Connectome-Based Predictive Modeling

We used connectome-based predictive modeling^23,39^ (CPM) to predict individual responses to all items of the experiential questionnaire, as well as the loadings on both axes of the low dimensional representation of in-scanner thoughts (i.e., TP1 and TP2). To mitigate the effect of head motion confounds, we residualized target variables (i.e., images score, TP1) with respect to head motion before training the models.

CPM is a technique that builds predictions of behavioral or clinical scores from whole-brain FC data. A detailed description of the method can be found in Shen et al.^39^. Here we will briefly describe the specifics of our implementation, which matches that of Finn & Bandettini^68^. First, data is divided into training and test sets using a 10-fold cross validation procedure. Second, in each training set, we looked for edges that significantly correlate (p< 0.01), positively or negatively, with the target of prediction. Next, those edges were used to construct the following model:

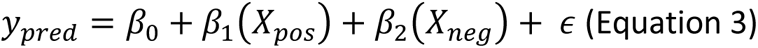

where *X*_*pos*_ and *X*_*neg*_ denote the sum of connectivity strength for all edges that correlated positively and negatively, respectively, with the prediction target. Finally, we used this model on the testing set to predict the target metric. These steps were conducted over the 10 different folds to gather predictions for all scans.

Once those were available, we evaluated prediction performance in terms of the Pearson’s correlation between observed and predicted values. It is worth noting that Pearson’s correlation is a relative indicator of predictive performance that affords us an understanding of how the model can distinguish between higher scores and lower scores. We emphasize that the results we report should be interpreted in this context as it is possible that models with the same correlation between observed and predicted values can have varying levels of absolute error (i.e., mean squared error).

We repeated this whole process 100 times to ensure our results were robust to variability in how the data was split into training and testing folds during the cross-validation step. Given that our dataset includes multiple scans per subject, it was important to consider subject identity during the split of the data into training and testing sets. When building folds, we ensured that all scans from the same subject were always included in the testing or training sets, but not across both. In other words, we did not use information from scans of one subject to predict a behavioral score for a different scan from the same subject.

##### Statistical Testing

To assess the statistical significance of the predictions, we generated a null distribution by running the whole CPM pipeline using randomized labels 10,000 times. We then computed a non-parametric p-value for the observed model accuracy using this formula^68^:

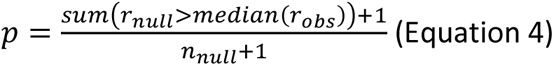

where *r*_*null*_ represents prediction accuracy values across all 10,000 null permutations, *median*(*r*_*obs*)_ is the median prediction accuracy across the 100 attempts made on a given target and *n*_*null*_ is the number of random permutations available (i.e., 10,000).

##### Predictive Model Visualization

In addition to the predicted scores, CPM also produces a network model of connections contributing to the prediction both in the positive direction (i.e., positive correlation between edge connectivity strength and predicted score) and negative direction (i.e., negative correlation between edge connectivity strength and predicted score). To generate these final models, we first calculated the average number of times a given edge contributed to a model across each of the 10 folds in each of the 100 iterations. We then only plotted and interpreted edges that were consistently included in the predictive models 90% of the time or more^68^.

#### Literature Search

##### Network-based Statistics Studies

Clinical studies that rely on Network-based Statistics (NBS) to search for significant differences in FC connectivity between clinical and healthy populations were identified as follows. We first used Scopus to find all open-access manuscripts that reference the original NBS work by Zalesky et al.^38^ and include the terms “functional MRI” (or variants) and “resting-state” (or variants) in the title, abstract or keyword list. The actual *Scopus* (http://scopus.com) query was:

*((REF(network-based AND statistic: AND identifying AND differences AND in AND brain AND networks) AND TITLE-ABS-KEY(( "functional MRI" OR "fMRI" OR "functional magnetic resonance imaging") AND ( "rest" OR "resting-state" OR "resting state" ))) AND PUBYEAR > 2012 AND PUBYEAR < 2024 AND ( LIMIT-TO ( DOCTYPE,"ar" ) ) AND ( LIMIT-TO ( LANGUAGE,"English" ) ) AND ( LIMIT-TO ( OA,"all" ) ) )*

This query returned a list of 402 documents. Next, going in reverse chronological order (oldest to newest), we search for the first 50 studies that report significant differences in resting-state static functional connectivity between healthy controls and a clinical population of interest based on some form of whole brain parcellation and that were conducted on human subjects. These criteria, set to match our study conceptual setup as much as possible, allowed us to discard search results with different study designs, such as pre-clinical studies, those based on task data, or dynamic aspects of connectivity, among many others.

For all 50 selected studies, we gathered—whenever available—basic information regarding sample size, scanning protocol (scanner strength, scan duration, repetition time), NBS hyper-parameters (edge-level threshold, number of permutations, network-level threshold), parcellation size, and number and percentage of connections (and edges) found to be significantly different. For studies that report multiple contrasts (e.g., more than one clinical condition or multi-site studies) we gathered information for all available contrasts. When results are reported at multiple edge-level thresholds, we gathered results for the threshold more similar to ours (i.e., T > 3.1). Suppl. Table 1 shows the studies that matched our criteria.

##### Connectome-Based Predictive Modeling Studies

Studies that apply connectome predictive modeling to resting-state fMRI data were identified as follows. We first used Scopus to find all open-access manuscripts that describe the use of connectome predictive modeling to resting-state data in the abstract and cite the original work of Shen et al.^39^ describing the CPM method, using the following Scopus query:

*( TITLE-ABS-KEY ( ( "resting-state" OR "resting state" ) AND ( "connectome predictive modeling" OR "connectome based predictive modeling") AND NOT dynamic AND NOT eeg AND NOT meg AND NOT fnirs ) AND REF ( using AND connectome-based AND predictive AND modeling AND to AND predict AND individual AND behavior AND from AND brain AND connectivity ) )*

This query returned a list of 88 documents. We then excluded from further analysis those studies that do not apply CPM to full brain ROI-based FC matrices (10 papers), use substantially modified versions of CPM (e.g., substitute OLS by SVR, have additional feature selection procedures, etc. | 8 papers), apply CPM only to task data (6 papers), report accuracy in a manner other than correlation between observed and predicted values (2 papers), report results for non-human data (1 paper), are based on non-fMRI data (1 paper), or are review manuscripts (3 papers). The remaining 57 studies reported accuracy for a total of 159 different CPM models, as several studies attempted prediction of more than one target and/or report accuracy separately for the positive, negative and combined CPM models. For studies that reported accuracy based on more than one internal cross validation scheme, we report that for 10-Fold cross validation whenever possible (as this is the one used here). Each model was then assigned a label based on the nature of the prediction target: Clinical (53 models), Cognition (32 models), Personality/Well-being (26 models).

Of the 159 CPM models selected and labeled, only 91 report accuracy in terms of the Pearson’s Correlation between observed and predicted values. It is these models that we use when constructing Figure 3d. The remaining models, which used Spearman’s correlation, were discarded when generating the figure.

Supplementary Table 2 provides the list of 57 studies that passed our exclusion criteria. We report title, year, parcellation, number of regions of interest, cross validation approach, and model accuracies. Supplementary Table 3 lists the studies found in Scopus that were excluded, as well as the exclusion criteria that applied to each particular study.

## Data Availability

This study is based on publicly available data as described in Mendes et al.^27^ Imaging data can be accessed at ftp://ftp.gwdg.de/pub/misc/MPILeipzig_Mind-Brain-Body. Behavioral data is available at https://dataverse.harvard.edu/dataset.xhtml?persistentId=doi:10.7910/DVN/VMJ6NV. Additional access locations can be found on the Mendes et al. publication.

## Code Availability

Code used in this study is available at https://github.com/nimh-sfim/fc_introspection

## Acknowledgements

This research was possible thanks to the support of the National Institute of Mental Health Intramural Research Programs (ZIAMH002783, ZICMH002968). Portions of this study used the high-performance computational capabilities of the Biowulf Linux cluster at the National Institutes of Health, Bethesda, MD (biowulf.nih.gov).

## Author contributions

Conceptualization: J.G.C., J.K. and P.B. Methodology: J.G.C., F.P., K.L., D.H. Software and resources: J.G.C., M.S., K.L. Formal analysis and visualization: J.G.G., M.S., I.G., K.L. Writing (original draft): J.G.C. Writing (review and editing): P.B., D.H., M.S., I.G., J.K., F.P., K.L.

