## Supplementary Materials for "In-Scanner Thoughts shape Resting-state Functional Connectivity: how participants “rest” matters"

### Supplementary Tables

| Year | Title | Clinical Condition | Healthy Controls (N) | Patients (N) | B0 | Dur(s) | TR(s) | Edge Threshold | Size | # Permut. | Network Threshold | # Nodes | # Edges | # Sig. Edges | % Sig. Edges | # Sig. Nodes | % Sig. Nodes |
|---|---|---|---|---|---|---|---|---|---|---|---|---|---|---|---|---|---|
| 2010 | Network-based statistic: Identifying differences in brain networks | Schizophrenia | 15 | 12 | 1.5T | 1024 | 2 | T > 3.0 | Extent | 5000 | pFWE = 0.037 | 74 | 2701 | 40 | 0.015 | 29 | 0.3919 |
| 2013 | Decreased Functional Brain Connectivity in Adolescents with Internet Addiction | Internet Addiction | 11 | 12 | 3.0T | 405 | 2.7 | T > 3.0 | Extent | 20000 | pFWE < 0.05 | 90 | 4005 | 59 | 0.015 | 38 | 0.422 |
| 2013 | Disrupted topological organization in whole-brain functional networks of heroin-dependent individuals: A resting-state fMRI study | Heroin Addiction | 15 | 17 | 1.5T | 480 | 2 | T > 1.7 | Extent | 1000 | pFWE < 0.001 | 90 | 4005 | 19 | 0.005 | 19 | 0.211 |
| 2014 | Altered brain network modules induce helplessness in major depressive disorder | Major Depressive Disorder | 16 | 16 | 3.0T | 300 | 3 | | Extent | 5000 | pFWE < 0.05 | 90 | 4005 | 24 | 0.006 | 24 | 0.267 |
| 2014 | Disruption of structure-function coupling in the schizophrenia connectome | Schizophrenia | 17 | 17 | 3.0T | 600 | 2.4 | T > 3.0 | Extent | 10000 | pFWE = 0.011 | 90 | 4005 | 86 | 0.021 | 72 | 0.8 |
| 2015 | Connectome-scale assessments of functional connectivity in children with primary monosymptomatic nocturnal enuresis | Nocturnal Enuresis | 29 | 24 | 3.0T | 420 | 2 | T > 1.6 | - | 10000 | pFWE < 0.05 | 90 | 4005 | 10 | 0.002 | 7 | 0.078 |
| 2015 | Disrupted brain network topology in pediatric posttraumatic stress disorder: A resting-state fMRI study | Posttraumatic Stress Disorder | 24 | 24 | 3.0T | 400 | 2 | T > 2.0 | - | 10000 | pFWE = 0.007 | 90 | 4005 | 7 | 0.002 | 13 | 0.144 |
| 2015 | Frequency dependent topological alterations of intrinsic functional connectome in major depressive disorder | Major Depressive Disorder | 50 | 50 | 3.0T | 484 | 2 | p <0.001 | Extent | 10000 | pFWE < 0.05 | 112 | 6216 | 23 | 0.004 | 13 | 0.116 |
| 2015 | Network topology and functional connectivity disturbances precede the onset of Huntington's disease | Huntington's Disease (medium disease burden) | 16 | 16 | 3.0T | 369.6 | 2.8 | T > 3.5 | - | 5000 | pFWE < 0.05 | 300 | 44850 | 96 | 0.002 | - | - |
| | | Huntington's disease (high disease burden) | | | | | | | | | | | | 252 | 0.006 | - | - |
| 2015 | Reduced functional connectivity in the thalamo-insular subnetwork in patients with acute anorexia nervosa | Anorexia Nervosa | 35 | 35 | 3.0T | 418 | 2.2 | T > 4.0 | Extent | 5000 | pFWE < 0.05 | 104 | 5356 | 7 | 0.001 | 7 | 0.067 |
| 2016 | Aberrant functional brain connectome in people with antisocial personality disorder | Antisocial Personality Disorder | 32 | 32 | 1.5T | 300 | 2 | p < 0.0005 | Extent | 10000 | pFWE = 0.001 | 90 | 4005 | 47 | 0.012 | 38 | 0.422 |
| 2016 | Altered functional connectivity of the default mode network in Williams syndrome: a multimodal approach | Williams Syndrome | 7 | 7 | 3.0T | 300 | 3 | p <0.01 | Extent | 5000 | pFWE = 0.027 | 82 | 3321 | 22 | 0.007 | 21 | 0.256 |

| | | | | | | | | | | | | | | | | | |
|---|---|---|---|---|---|---|---|---|---|---|---|---|---|---|---|---|---|
| 2016 | Brain Connectomics' Modification to Clarify Motor and Nonmotor Features of Myotonic Dystrophy Type 1 | Myotonic Dystrophy Type 1 | 26 | 31 | 3.0T | 440 | 2.08 | | - | 50000 | pFWE < 0.005 | 116 | 6670 | 83 | 0.012 | 51 | 0.44 |
| 2016 | Disrupted topological organization of structural and functional brain connectomes in clinically isolated syndrome and multiple sclerosis | Multiple Sclerosis | 35 | 41 | 3.0T | 360 | 2 | p < 0.05 | Extent | 10000 | pFWE < 0.05 | 90 | 4005 | 38 | 0 | 22 | 0 |
| 2016 | Distinct disruptions of resting-state functional brain networks in familial and sporadic schizophrenia | Schizophrenia (familiar) | 26 | 26 | 3.0T | 370 | 2 | T > 2.01 | - | 10000 | pFWE = 0.022 | 90 | 4005 | 13 | 0.003 | 13 | 0.144 |
| | | Schizophrenia (sporadic) | | | | | | | | | pFWE = 0.033 | | | 12 | 0.003 | 12 | 0.133 |
| 2016 | Large-scale hypoconnectivity between resting-state functional networks in unmedicated adolescent major depressive disorder | Major Depressive Disorder | 56 | 55 | 3.0T | 512 | 2 | T > 1.659, p < 0.05 | Extent | 100000 | pFWE=0.0237 | 17 | 136 | 7 | 0.051 | 8 | 0.471 |
| 2016 | Network analysis of functional brain connectivity in borderline personality disorder using resting-state fMRI | Borderline Personality Disorder | 10 | 20 | 3.0T | 360 | 2 | T > 3.5 | Extent | 10000 | pFWE = 0.0304 | 82 | 3321 | 26 | 0.017 | 26 | 0.488 |
| 2016 | Resting state brain network disturbances related to hypomania and depression in medication-free bipolar disorder | Bipolar Disorder | 30 | 60 | 3.0T | 333 | 2.25 | T > 3.35 | Extent | 5000 | pFWE < 0.05 | 181 | 16290 | 61 | 0.004 | 48 | 0.2652 |
| 2016 | Whole-brain analytic measures of network communication reveal increased structure-function correlation in right temporal lobe epilepsy | Epilepsy | 13 | 7 | 3.0T | 1200 | 3.6 | T > 5.0 | Extent | 10000 | pFWE < 0.05 | 512 | 130816 | 104 | 0.001 | 38 | 0.074 |
| 2017 | Altered functional network architecture in orbitofronto-striato-thalamic circuit of unmedicated patients with obsessive-compulsive disorder | Obsessive Compulsive Disorder | 61 | 61 | 3.0T | 413 | 3.5 | p < 0.05 | - | 10000 | pFWE < 0.05 | 17 | 136 | 8 | 0.059 | 10 | 0.5882 |
| 2017 | Altered intrinsic functional brain architecture in female patients with bulimia nervosa | Bulimia Nervosa | 45 | 48 | 3.0T | 420 | 2 | | - | 10000 | - | 90 | 4005 | 103 | 0.026 | 21 | 0.2333 |
| 2017 | Altered network efficiency of functional brain networks in patients with breast cancer after chemotherapy | Breast Cancer | 40 | 28 | - | - | - | p < 001 | Extent | 10000 | - | 90 | 4005 | 79 | 0.02 | 35 | 0.3889 |
| 2017 | Cognitive phenotypes in parkinson's disease differ in terms of brain-network organization and connectivity | Parkinson's Disease (G1 > G2) | 31 | 32 | 3.0T | 600 | 2.4 | p < 0.005 | Extent | 20000 | - | 164 | 13366 | 19 | 0.001 | 20 | 0.122 |
| | | Parkinson's Disease (G1 > G3) | | 43 | | | | | | | | | | 63 | 0.005 | 63 | 0.3841 |
| 2017 | Cognitive phenotypes in parkinson's disease differ in terms of brain-network organization and connectivity | Parkinson's Disease (G1 > G4) | 31 | 14 | 3.0T | 600 | 2.4 | p < 0.005 | Extent | 20000 | - | 164 | 13366 | 172 | 0.013 | 128 | 0.7805 |
| 2017 | Decreased functional connectivity within a language subnetwork in benign epilepsy with centrotemporal spikes | Epilepsy | 20 | 25 | 3.0T | 368 | 2 | T > 3.0 | Intensity | 50000 | - | 90 | 4005 | 41 | 0.01 | 36 | 0.4 |
| 2017 | Discriminating cognitive status in Parkinson's disease through functional connectomics and machine learning | Parkinson's Disease | 38 | 27 | 3.0T | 600 | 2 | - | Extent | 10000 | - | 246 | 30135 | 235 | 0.008 | 120 | 0.4878 |

| | | | | | | | | | | | | | | | | | |
|---|---|---|---|---|---|---|---|---|---|---|---|---|---|---|---|---|---|
| 2017 | Disrupted functional connectome in antisocial personality disorder | Antisocial Personality Disorder | 32 | 32 | 1.5T | 300 | 2 | p < 5e-4 | Extent | | - | 90 | 4005 | 47 | 0.012 | 38 | 0.4222 |
| 2017 | Disrupted resting-state brain network properties in obesity: Decreased global and putaminal cortico-striatal network efficiency | Obesity | 40 | 40 | 3.0T | | | T > 3.0 | Extent | 10000 | - | 90 | 4005 | 30 | 0.007 | 19 | 0.2111 |
| 2017 | Evaluation of Whole-Brain Resting-State Functional Connectivity in Spinal Cord Injury: A Large-Scale Network Analysis Using Network-Based Statistic | Spinal Cord Injury | 15 | 15 | 3.0T | 480 | 2 | - | - | - | - | 116 | 6670 | 21 | 0.003 | 20 | 0.1724 |
| 2017 | Neural substrates of motor and cognitive dysfunctions in SCA2 patients: A network based statistics analysis | Spinocerebellar Ataxia Type 2 | 33 | 9 | 3.0T | 440 | 2.08 | - | - | 5000 | pFWE < 0.05 | 116 | 6670 | 110 | 0.016 | 62 | 0.5345 |
| 2017 | Reduced orbitofrontal-thalamic functional connectivity related to suicidal ideation in patients with major depressive disorder | Major Depressive Disorder | 36 | 36 | 3.0T | 360 | 3 | T > 3.0 | Extent | 5000 | pFWE < 0.05 | 90 | 4005 | 6 | 0.001 | 7 | 0.0778 |
| 2017 | Structural and functional hyperconnectivity within the sensorimotor system in xenomelia | Xenomelia | 13 | 13 | 3.0T | 600 | 4 | T > 3.6 | Extent | 5000 | - | 116 | 6670 | 24 | 0.004 | 27 | 0.2328 |
| 2017 | Tackling variability: A multicenter study to provide a gold-standard network approach for frontotemporal dementia | Fronto-Temporal Dementia (Country 1) | 16 | 16 | 1.5T | 600 | 2.7 | T > 3.0 | Extent | 5000 | - | 90 | 4005 | 51 | 0.013 | 42 | 0.4667 |
| | | Fronto-Temporal Dementia (Country 2) | 29 | 17 | 3.0T | 300 | 3 | T > 3.0 | | | | | | 83 | 0.021 | 56 | 0.6222 |
| | | Fronto-Temporal Dementia (Country 3) | 15 | 12 | 3.0T | 420 | 2 | T > 4.0 | | | | | | 40 | 0.01 | 35 | 0.3889 |
| 2017 | Topologically convergent and divergent functional connectivity patterns in unmedicated unipolar depression and bipolar disorder | Unipolar Depression | 31 | 43 | - | - | - | p < 0.001 | Extent | 5000 | - | 1024 | 523776 | 51 | 0 | 44 | 0.043 |
| | | Bipolar Disorder | 32 | | | | | | | | | | | 81 | 0 | 74 | 0.0723 |
| 2018 | Altered brain functional connectome in migraine with and without restless legs syndrome: A resting-state functional MRI study | Migraine w/o RLS | 19 | 22 | 3.0T | 500 | 2.5 | p < 0.005 | Extent | 5000 | pFWE < 0.05 | 236 | 27730 | 162 | 0.006 | 121 | 0.5127 |
| | | Migraine with RLS | | | | | | | | | | | | 136 | 0.005 | 82 | 0.3475 |
| 2018 | Altered cerebellar-insular-parietal-cingular subnetwork in adolescents in the earliest stages of anorexia nervosa: A network-based statistic analysis | Anorexia Nervosa | 15 | 15 | 1.5T | 284.8 | 3.56 | T > 3.1 | - | 10000 | pFWE < 0.05 | 128 | 8128 | 31 | 0.004 | 30 | 0.2344 |
| 2018 | Brain-behavior patterns define a dimensional biotype in medication-naïve adults with attention-deficit hyperactivity disorder | Attention Deficit Hyperactivity Disorder | 123 | 80 | 3.0T | 459 | 2.55 | T > 3.5 | - | 10000 | pFWE < 0.05 | 253 | 31878 | 41 | 0.001 | 40 | 0.1581 |
| 2018 | Evaluating functional connectivity alterations in autism spectrum disorder using network-based statistics | Autism Spectrum Disorder | 82 | 74 | 3.0T | 360 | 2 | T > 3.0 | Extent | 5000 | pFWE < 0.025 | 116 | 6670 | 158 | 0.024 | 105 | 0.9052 |

| | | | | | | | | | | | | | | | | | |
|---|---|---|---|---|---|---|---|---|---|---|---|---|---|---|---|---|---|
| 2018 | Impact of Zika Virus on adult human brain structure and functional organization | Zika Virus | 9 | 9 | 3.0T | 360 | 2 | T > 3.6 | Extent | 5000 | - | 160 | 12720 | 19 | 0.001 | 20 | 0.125 |
| 2018 | Progressively disrupted brain functional connectivity network in subcortical ischemic vascular cognitive impairment patients | Subcortical Ischemic Vascular Cognitive Impairment w/o Dementia | 19 | 20 | 3.0T | 480 | 2 | p < 0.005 | Extent | 10000 | - | 90 | 4005 | 93 | 0.023 | 38 | 0.4222 |
| | | Subcortical Ischemic Vascular Cognitive Impairment with Dementia | | | | | | | | | | | | 141 | 0.035 | 66 | 0.7333 |
| 2018 | Short- and long-range synergism disorders in lifelong premature ejaculation evaluated using the functional connectivity density and network property | Alcohol Dependent | 36 | 21 | 3.0T | 360 | 3 | T > 3.0 | Extent | 10000 | pFWE < 0.05 | 90 | 4005 | 373 | 0.093 | 69 | 0.7667 |
| 2019 | Aberrant brain network topology in fronto-limbic circuitry differentiates euthymic bipolar disorder from recurrent major depressive disorder | Bipolar Disorder | 30 | 20 | 3.0T | 600 | 2 | T > 3.4 | Intensity | 10000 | pFWE < 0.05 | 90 | 4005 | 11 | 0.003 | 11 | 0.1222 |
| | | | 42 | 57 | | 480 | | T > 3.1 | Extent | | | | | 85 | 0.021 | 68 | 0.7556 |
| 2019 | Brain functional connectivity is altered in patients with Takotsubo Syndrome | Takotsubo Syndrome | 8 | 8 | 1.5T | 380 | 2 | T > 4.14 | | 5000 | pFWE < 0.05 | 268 | 35778 | 39 | 0.001 | 36 | 0.1343 |
| 2019 | Cognitive performance in mid-stage Parkinson's disease: functional connectivity under chronic antiparkinson treatment | Parkinson's Disease | 16 | 16 | 3.0T | | 2 | T > 6.0 | Intensity | 5000 | pFWE < 0.05 | 206 | 21115 | 15 | 0.001 | 20 | 0.0971 |
| 2019 | Data-Driven Clustering Reveals a Link Between Symptoms and Functional Brain Connectivity in Depression | Depression | 72 | 178 | 3.0T | 500 | 2.5 | p < 0.05 | | 10000 | pFWE < 0.05 | 19 | 171 | 22 | 0.129 | 15 | 0.7895 |
| 2019 | Disrupted topological organization of human brain connectome in diabetic retinopathy patients | Diabetic Retinopathy | 38 | 35 | 3.0T | | 2 | T > 2.649 | - | 10000 | - | 90 | 4005 | 52 | 0.013 | 40 | 0.4444 |
| 2019 | Early functional connectivity predicts recovery from visual field defects after stroke | Stroke | 15 | 32 | 3.0T | 240 | 2 | T > 3.0 | Extent | 5000 | pFWE < 0.05 | 82 | 3321 | 9 | 0.003 | 6 | 0.0732 |
| 2019 | Functional brain connectome and its relation to mild cognitive impairment in cerebral small vessel disease patients with thalamus lacunes: A cross-sectional study | Cerebral Small Vessel Disease with Thalamus Lacunes | 34 | 14 | 3.0T | 420 | 2 | T > 2.02 | - | 10000 | - | 90 | 4005 | 13 | 0.003 | 6 | 0.0667 |
| 2019 | Functional network-based statistics reveal abnormal resting-state functional connectivity in minimal hepatic encephalopathy | Anorexia Nervosa | 19 | 19 | 3.0T | 360 | 2 | T > 3.1 | - | 5000 | pFWE < 0.05 | 90 | 4005 | 43 | 0.011 | 29 | 0.3222 |
| 2019 | Resting-state functional connectivity after concussion is associated with clinical recovery | Concussion | 60 | 62 | 3.0T | 360.72 | 0.72 | T > 2.5 | - | 10000 | pFWE < 0.05 | 200 | 19900 | 1464 | 0.074 | 180 | 0.9 |

**Supplementary Table 1 |** Studies selected for literature review of prior NBS-based clinical studies in human populations.

#### Supplementary Table 2.

| Year | Title | Category | Prediction Target | # Subjects | Atlas | # ROIs | Model Accuracy (r) | | | | | | Cross Validation |
|---|---|---|---|---|---|---|---|---|---|---|---|---|---|
| | | | | | | | Positive Network | | Negative Network | | Pos + Neg Networks | | |
| | | | | | | | Pearson | Spearman | Pearson | Spearman | Pearson | Spearman | |
| 2018 | Individualized prediction of trait narcissism from whole-brain resting-state functional connectivity | Personality | Narcissism | 168 | Shen 268-node | 268 | **0.24** | - | - | - | - | - | LOOCV |
| 2018 | Resting-state functional connectivity predicts cognitive impairment related to Alzheimer's disease | Clinical | Alzheimer's Disease Assessment Scale (ADAS 11) Score | 59 | Shen 268-node | 268 | - | **0.49** | - | 0.27 | - | - | LOOCV |
| 2018 | Resting-state functional connectivity predicts neuroticism and extraversion in novel individuals | Personality | Neuroticism | 114 | Shen 268-node | 244 | **0.27** | - | 0.14 | - | **0.22** | - | LOOCV |
| | | | Extraversion | | | | **0.22** | - | 0.2 | - | **0.22** | - | |
| 2019 | Connectome-based individualized prediction of loneliness | Cognitive | Loneliness | 75 | Shen 268-node | 268 | -0.3 | - | **0.244** | - | - | - | LOOCV |
| 2019 | The Functional Brain Organization of an Individual Allows Prediction of Measures of Social Abilities Transdiagnostically in Autism and Attention-Deficit/Hyperactivity Disorder | Cognition | Processing Speed | 99 | Shen 268-node | 268 | - | **0.35** | - | **0.4** | - | **0.38** | 10-Fold CV |
| | | Clinical | Social Responsiveness Scale (SRS) Total Score | | | | - | - | - | - | **0.32** | - | LOOCV |
| | | | SRS Communication score | | | | - | - | - | - | **0.3** | - | |
| | | | SRS Motivation score | | | | - | - | - | - | **0.23** | - | |
| | | | SRS Mannerisms score | | | | - | - | - | - | **0.37** | - | |
| | | | SRS Awareness score | | | | - | - | - | - | **0.27** | - | |
| | | | SRS Cognition score | | | | - | - | - | - | **0.27** | - | |
| | | | Autism Diagnostic Observation Scale (ADOS) Total score | | | | - | - | - | - | **0.43** | - | |
| | | | Ados Social Affect Score | | | | - | - | - | - | **0.53** | - | |
| | | | Ados Generic Total Score | | | | - | - | - | - | **0.4** | - | |
| | | | Ados Severity Score | | | | - | - | - | - | **0.6** | - | |
| 2020 | Behavioral and brain signatures of substance use vulnerability in childhood | Clinical | Substance Risk Seeking | 3193 | Shen 268-node | 268 | - | - | - | - | **0.1** | - | 10-Fold CV |
| | | | Familial Risk For Substance Abuse | 3193 | | | - | - | - | - | - | - | |

| | | | | | | | | | | | | | |
|---|---|---|---|---|---|---|---|---|---|---|---|---|---|
| 2020 | Brain functional connectome-based prediction of individual decision impulsivity | Personality | Decision Impulsivity (Ddis_AUC_40K) | 809 | Group-ICA | 200 | **0.25** | - | **0.24** | - | - | - | LOOCV |
| | | | Decision Impulsivity (Ddis_AUC_200) | | | | **0.23** | - | 0.19 | - | - | - | |
| 2020 | Connectome-based models can predict early symptom improvement in major depressive disorder | Clinical | Depression Severity | 108 | Shen 268-node | 268 | - | - | **0.41** | - | - | - | LOOCV |
| 2020 | Connectome-based models can predict processing speed in older adults | Cognitive | Processing Speed | 99 | Shen 268-node | 268 | - | **0.36** | - | **0.42** | - | **0.4** | LOOCV |
| 2020 | Do intrinsic brain functional networks predict working memory from childhood to adulthood? | Cognitive | Working Memory | 87 - Preschoolers | Shen 268-node | 268 | - | - | - | - | - | 0.118 | 5-Fold CV |
| | | | | 117 - Early school age | | | - | - | - | - | - | 0.04 | |
| | | | | 110 - Late school age | | | - | - | - | - | - | -0.009 | |
| | | | | 95 - Adolescents | | | - | - | - | - | - | **0.228** | |
| | | | | 59 - Adults | | | - | - | - | - | - | **0.312** | |
| 2020 | Individualized Prediction of PTSD Symptom Severity in Trauma Survivors From Whole-Brain Resting-State Functional Connectivity | Clinical | PTSD Symptoms | 64 | AAL | 268 | - | **0.3** | - | **0.17** | - | - | LOOCV |
| 2020 | Multimodal data revealed different neurobiological correlates of intelligence between males and females | Cognitive | Intelligence Quotient | 166 - Males | AAL | 116 | **0.22** | - | **0.27** | - | **0.28** | - | LOOCV |
| | | | | 160 - Females | | | **0.23** | - | **0.33** | - | **0.36** | - | |
| 2020 | Predicting Patient Reported Outcomes of Cognitive Function Using Connectome-Based Predictive Modeling in Breast Cancer | Clinical | Executive Dysfunction | 76 | Shen 268-node | 268 | - | - | - | - | **0.65** | - | 10-Fold CV |
| | | | Memory Functions | | | | - | - | - | - | **0.32** | - | |
| 2020 | Preliminary prediction of individual response to electroconvulsive therapy using whole-brain functional magnetic resonance imaging data | Clinical | Change In Depression Scores | 122 | Brainnetome Atlas | 246 | - | **0.27** | - | **0.51** | - | **0.46** | LOOCV |
| 2020 | Robust prediction of individual personality from brain functional connectome | Personality | Agreeableness | 1003 | ICA derived nodes | 200 | **0.163** | - | 0.115 | - | - | - | LOOCV |
| | | | Openness | | | | -0.135 | - | 0.138 | - | - | - | |
| | | | Conscientiousness | | | | 0.124 | - | **0.281** | - | - | - | |
| | | | Neuroticism | | | | 0.097 | - | -0.085 | - | - | - | |
| | | | Extraversion | | | | -0.037 | - | -0.078 | - | - | - | |
| 2020 | | Cognitive | Digit Span Forward | 91 | AAL | 90 | 0.083 | - | **0.51** | - | **0.25** | - | LOOCV |

| | | | | | | | | | | | | | |
|---|---|---|---|---|---|---|---|---|---|---|---|---|---|
| | The individualized prediction of cognitive test scores in mild cognitive impairment using structural and functional connectivity features | | Digit Span Backward | | | | **0.49** | - | **0.6** | - | **0.59** | - | |
| | | | Immediate Recall | | | | **0.38** | - | **0.54** | - | **0.51** | - | |
| | | | Delayed Recall | | | | **0.35** | - | **0.46** | - | **0.43** | - | |
| | | | Recognition | | | | **0.33** | - | 0.099 | - | **0.31** | - | |
| | | | Color Trails Part 1 | | | | **0.55** | - | **0.58** | - | **0.56** | - | |
| | | | Color Trails Part 2 | | | | **0.6** | - | **0.65** | - | **0.63** | - | |
| | | | Color Trains Test Inference | | | | 0.11 | - | **0.41** | - | 0.21 | - | |
| | | | Wais-Iii Block Design | | | | **0.55** | - | **0.45** | - | **0.48** | - | |
| | | | Mini-Mental State Examination | | | | 0.51 | - | **0.52** | - | **0.54** | - | |
| 2021 | Connectome-based model predicts episodic memory performance in individuals with subjective cognitive decline and amnestic mild cognitive impairment | Cognition | Auditory/Verbal Test Delayed Call | 83 | AAL | 116 | 0.033 | - | **0.22** | - | - | - | LOOCV |
| 2021 | Connectome-based prediction of global cognitive performance in people with HIV | Clinical | Global Cognitive Measure | 67 | Shen 268-node | 268 | **0.35** | - | **0.382** | - | **0.464** | - | LOOCV |
| 2021 | Connectome-Based Predictive Modeling of Creativity Anxiety | Clinical | Creativity Anxiety | 281 | Shen 268-node | 268 | **0.18** | - | **0.2** | - | **0.19** | - | LOOCV |
| 2021 | Connectome-Based Predictive Modeling of Individual Anxiety | Clinical | Anxiety | 76 | Shen 268-node | 268 | - | -0.01 | - | **0.33** | - | - | LOOCV |
| 2021 | The functional and structural connectomes associated with geriatric depression and anxiety symptoms in mild cognitive impairment: Cross-syndrome overlap and generalization | Clinical | Geriatric Depression Scale (GDS) | 91 | Brainnetome atlas | 246 | - | **0.331** | - | **0.314** | - | **0.357** | LOOCV |
| | | | Geriatric Anxiety Inventory (GAI) | | | | - | **0.198** | - | **0.308** | - | **0.308** | |
| 2021 | The functional connectome predicts feeling of stress on regular days and during the COVID-19 pandemic | | Perceived Stress | 673 | Brainnetome Atlas | 246 | **0.36** | - | **0.41** | - | **0.44** | - | LOOCV |
| 2022 | Connectome-based model predicts individual psychopathic traits in college students | Personality | T-PTS (Psychopathic traits) | 84 | Dosenbach | 142 | - | 0.073 | - | **0.295** | - | **0.308** | LOOCV and k fold |
| | | | S-PTS (Psychopathic traits) | | | | - | 0.129 | - | **0.309** | - | **0.353** | |
| | | | P-PTS (Psychopathic traits) | | | | - | 0.008 | - | 0.07 | - | 0.099 | |
| 2022 | Connectome-based prediction of marital quality in husbands' processing of spousal interactions | Well-being | Marital Quality | 25 - Males | Shen 268-node | 268 | -0.15 | - | -0.08 | - | -0.31 | - | LOOCV |
| | | | | 25 - Females | | | -0.39 | - | 0.17 | - | -0.07 | - | |

| | | | | | | | | | | | | | |
|---|---|---|---|---|---|---|---|---|---|---|---|---|---|
| 2022 | Connectome-based predictive models using resting-state fMRI for studying brain aging | Demographic | Age | 256 | Shen 268-node | 268 | **0.398** | - | **0.505** | - | **0.484** | - | LOOCV |
| 2022 | Individualized prediction of consummatory anhedonia from functional connectome in major depressive disorder | Clinical | Anhedonia | 54 | Shen 268-node | 268 | **0.28** | - | - | - | - | - | 10-Fold CV |
| 2022 | Multi-modality connectome-based predictive modeling of individualized compulsions in obsessive-compulsive disorder | Clinical | Compulsion | 54 | Shen 268-node | 268 | - | - | **0.33** | - | - | - | 10-Fold CV |
| | | | YBOCS Total Score | | | | - | - | **0.38** | - | **0.41** | - | LOOCV |
| 2022 | Neural mechanisms underlying empathy during alcohol abstinence: evidence from connectome-based predictive modeling | Personality | Empathy | 59 | Dosenbach | 160 | - | - | - | - | **0.24** | - | LOOCV |
| 2022 | Predicting children's math skills from task-based and resting-state functional brain connectivity | Cognition | Math Skills | 31 | Brainnetome Atlas | 210 | - | - | - | - | **0.54** | - | LOOCV |
| 2022 | Preoperative brain connectome predicts postoperative changes in processing speed in moyamoya disease | Clinical | Post-Operative Processing Speed (1 Month) | 12 | AAL | 116 | - | **0.63** | - | 0.31 | - | - | LOOCV |
| | | | Post-Operative Processing Speed (6 Month) | | | | - | **0.62** | - | **0.55** | - | - | |
| 2023 | A connectome-based neuromarker of nonverbal number acuity and arithmetic skills | Cognitive | Numeracy | 154 | Shen 268-node | 268 | **0.24** | - | 0.032 | - | - | - | LOOCV |
| 2023 | Age-related intrinsic functional connectivity underlying emotion utilization | Personality | Emotion Utilization | 133 | Shen 268-node | 219 | - | **0.29** | - | **0.2** | - | - | LOOCV |
| 2023 | Brain connectivity in frailty: Insights from The Irish Longitudinal Study on Ageing (TILDA) | | Frailty | 347 | Shen 268-node | 268 | - | **0.169** | - | **0.195** | - | - | 10-Fold CV |
| 2023 | Brief intensive social gaze training reorganizes functional brain connectivity in boys with fragile X syndrome | Clinical | FXS vs. ASD | 37 | Corgon Atlas | 333 | - | **0.37** | - | **0.44** | - | - | LOOCV |
| 2023 | Connectome-based fingerprint of motor impairment is stable along the course of Parkinson's disease | Clinical | Levels Of Motor Impairment | 81 | Shen 268-node | 268 | - | **0.21** | - | 0.16 | - | - | LOOCV |
| 2023 | Connectome-based modeling reveals a resting-state functional network that mediates the relationship between social rejection and rumination | Cognitive | Rumination | 560 | 400 ROI Schaefer | 400 | 0.045 | - | **0.152** | - | - | - | LOOCV |

| | | | | | | | | | | | | | |
|---|---|---|---|---|---|---|---|---|---|---|---|---|---|
| 2023 | Connectome-based prediction of eating disorder-associated symptomatology | Clinical | Body Image Concerns | 660 | Shen 268-node | 268 | - | **0.11** | - | **0.11** | - | **0.14** | 10-Fold CV |
| | | | Binge Eating Frequency | | | | - | 0.06 | - | 0.01 | - | 0.04 | |
| | | | Compensatory Behaviors | | | | - | 0.07 | - | **0.09** | - | **0.1** | |
| 2023 | Connectome-based prediction of the severity of autism spectrum disorder | Clinical | Communication | 359 - 323 ASD, 36 controls | Dosenbach Atlas | 160 | - | -0.18 | - | **0.22** | - | - | LOOCV |
| | | | Social Interaction | 359 - 323 ASD, 36 controls | | | - | **0.21** | - | 0.12 | - | - | |
| | | | Stereotyped Behavior | 359 - 323 ASD, 36 controls | | | - | -0.02 | - | 0.12 | - | - | |
| 2023 | Connectome-based predictive modeling of compulsion in obsessive-compulsive disorder | Clinical | Compulsion | 57 | Shen 268-node | 268 | - | **0.3** | - | **0.32** | - | 0.3 | LOOCV |
| 2023 | Connectome-based predictive modeling of fluid intelligence: evidence for a global system of functionally integrated brain networks | Cognitive | Fluid Intelligence | 159 | Glasser Atlas | - | **0.42** | - | - | - | - | - | LOOCV |
| 2023 | Connectome-based predictive modeling of trait forgiveness | Personality | Trait Forgiveness | 100 | Shen 268-node | 268 | **0.23** | - | **0.25** | - | **0.31** | - | 10-Fold CV |
| 2023 | Connectome-based predictive modeling predicts paranoid ideation in young men with paranoid personality disorder: a resting-state functional magnetic resonance imaging study | Clinical | Paranoia | 18 | Brainnetome Atlas | 246 | - | −0.0926 | - | **0.684** | - | 0.3148 | LOOCV |
| 2023 | Functional brain connectivity predicts sleep duration in youth and adults | Cognitive | Sleep Duration | 652 - HCP | Shen 268-node | 268 | - | - | - | - | - | **0.164** | 10-Fold CV |
| | | | | 786 - ABCD | | | - | - | - | - | - | **0.238** | |
| 2023 | Functional connectome predicting individual gait function and its relationship with molecular architecture in Parkinson's disease | Clinical | Gait Function | 48 | Shen 268-node | 268 | - | - | - | - | **0.31** | - | LOOCV |
| 2023 | Functional connectomes of akinetic-rigid and tremor within drug-naïve Parkinson's disease | Clinical | AR | 81 | Shen 268-node | 268 | - | - | - | - | - | **0.28** | LOOCV |
| | | | Tremor | | | | - | - | - | - | - | **0.32** | |
| 2023 | Neural correlates of schizotypal traits: Findings from connectome-based predictive modelling | Clinical | Schizophrenia Traits | 82 | Shen 268-node | 268 | -0.23 | - | **0.29** | - | - | - | LOOCV |
| 2023 | Task and Resting-State Functional Connectivity Predict Driving Violations | Personality | Sensation Seeking | 29 | Shen 268-node | 178 | 0.05 | - | -0.03 | - | - | - | LOOCV |
| | | | Impulsivity | | | | -0.27 | - | -0.09 | - | - | - | |
| | | | Driving Lapses | | | | **0.4** | - | -0.56 | - | - | - | |
| | | | Driving Errors | | | | 0.12 | - | **0.24** | - | - | - | |

| | | | | | | | | | | | | | |
|---|---|---|---|---|---|---|---|---|---|---|---|---|---|
| | | | Driving Violations | | | | -0.26 | - | -0.63 | - | - | - | |
| 2023 | The brain network underlying attentional blink predicts symptoms of attention deficit hyperactivity disorder in children | Clinical | ADHD Symptoms - AB | 91 | Shen 268-node | 268 | **0.284** | - | 0.395 | - | - | - | LOOCV |
| | | | ADHD Symptoms - TD | | | | **0.326** | - | **0.4** | - | - | - | |
| 2023 | The connectome-based prediction of trust propensity in older adults: A resting-state functional magnetic resonance imaging study | Personality | Trust Propensity | 120 | Dosenbach | 142 | - | **0.25** | - | 0.05 | - | - | 10-Fold CV |
| 2023 | The Individualized Prediction of Neurocognitive Function in People Living With HIV Based on Clinical and Multimodal Connectome Data | Cognitive | Cognitive Function (7 Domains) | 102 | Power Atlas | 264 | **0.35** | - | **0.45** | - | **0.54** | - | LOOCV |
| 2023 | Trait repetitive negative thinking in depression is associated with functional connectivity in negative thinking state rather than resting state | Clinical | MDD | 62 | Shen 268-node | 230 | - | - | - | - | - | **0.826** | 5-Fold CV |
| | | | Ruminative Response Scale | 36 - MDD | | | - | - | - | - | - | -0.049 | |
| | | | Ruminative Response Scale | 62 | | | - | - | - | - | - | **0.613** | |
| 2023 | Using modular connectome-based predictive modeling to reveal brain-behavior relationships of individual differences in working memory | Cognitive | Working memory (0-back d') | 874 | Shen 268-node | 268 | - | - | - | - | **0.18** | - | LOOCV |
| | | | Working memory (2-back d') | | | | - | - | - | - | **0.18** | - | |
| 2024 | Connectome-based prediction of decreased trust propensity in older adults with mild cognitive impairment: A resting-state functional magnetic resonance imaging study | Personality | Trust Propensity | 115 | Dosenbach | 142 | **0.41** | - | **0.2** | - | - | - | LOOCV |
| 2024 | Connectome-based predictive modeling of Internet addiction symptomatology | Clinical | Internet Addiction | 677 | Shen 268-node | 268 | -0.06 | - | **0.202** | - | - | - | LOOCV |
| 2024 | Connectome-based predictive modelling estimates individual cognitive status in Parkinson's disease | Clinical | Global Cognitive Composite Score | 58 | 368 Shen Atlas | 368 | - | **0.68** | - | **0.63** | - | **0.59** | LOOCV |
| 2024 | Finger motor representation supports the autonomy in arithmetic: neuroimaging evidence from abacus training | Cognitive | Arithmetic Ability | 69 - Control | 400 ROI Schaefer | 400 | - | **0.352** | - | 0.236 | - | - | LOSOCV (subject) |
| | | | | 78 - AMC (received math training prior) | | | - | **0.391** | - | **0.371** | - | - | |
| 2024 | Sex differences in functional connectivity and the predictive role of the connectome-based predictive model in Alzheimer's disease | Clinical | Montreal Cognitive Assessment Scores | 71 - Males | AAL | 90 | **0.23** | - | - | - | - | - | LOOCV |
| | | | | 88 - Females | | | **0.34** | - | - | - | - | - | |

**Supplementary Table 2 |** Studies selected for literature review of prior studies that use Connectome Predictive Modeling on resting-state fMRI. The 57 papers that passed our exclusion criteria with their respective year, descriptive category, target for prediction, number of subjects, atlas, number of regions of interest, cross validation method, and the model accuracies reported as either Pearson's or Spearman's R. Papers are listed alphabetically, by year. Bolded R values were reported as significant in the original papers. The papers which used Pearson correlation were used to generate Figure 3.d.

*This study reported the CPM accuracy before and after accounting for head motion, age, gender, and intelligence. We report the accuracy after accounting for these confounds, as we account for head motion in our analysis.

**Supplementary Table 3.**

| Year | Title | Exclusion Criteria |
|---|---|---|
| 2018 | Connectome-based models predict separable components of attention in novel individuals | CPM on task data |
| 2018 | Connectome-based predictive modeling of attention: Comparing different functional connectivity features and prediction methods across datasets | Review Article |
| 2018 | Robust prediction of individual creative ability from brain functional connectivity | CPM on task data |
| 2019 | An information network flow approach for measuring functional connectivity and predicting behavior | Substantially modified CPM |
| 2019 | Combining multiple connectomes improves predictive modeling of phenotypic measures | Review Article |
| 2019 | Connectome-based model predicts individual differences in propensity to trust | Substantially modified CPM |
| 2020 | Bootstrapping promotes the RSFC-behavior associations: An application of individual cognitive traits prediction | Incompatible Accuracy Reporting |
| 2020 | Functional connectomes linking child-parent relationships with psychological problems in adolescence | Not Whole Brain / ROI Level |
| 2020 | Individual variability in functional connectivity architecture of the mouse brain | Non Human data |
| 2021 | Context Matters: Situational Stress Impedes Functional Reorganization of Intrinsic Brain Connectivity during Problem-Solving | EEG Study |
| 2021 | Dissociable neural substrates of opioid and cocaine use identified via connectome-based modelling | CPM on task data |
| 2021 | Impact of binge drinking during college on resting state functional connectivity | Not Whole Brain / ROI Level |
| 2021 | NBS-Predict: A prediction-based extension of the network-based statistic | Substantially modified CPM |
| 2021 | Neurobiological substrates of the positive formal thought disorder in schizophrenia revealed by seed connectome-based predictive modeling | Not Whole Brain / ROI Level |
| 2021 | Optimizing differential identifiability improves connectome predictive modeling of cognitive deficits from functional connectivity in Alzheimer's disease | Substantially modified CPM |
| 2021 | Resting-state connectome-based support-vector-machine predictive modeling of internet gaming disorder | Substantially modified CPM |
| 2021 | Time-delay structure predicts clinical scores for patients with disorders of consciousness using resting-state fMRI | Substantially modified CPM |
| 2022 | Functional connectivity of the central autonomic and default mode networks represent neural correlates and predictors of individual personality | Not Whole Brain / ROI Level |
| 2022 | Functional Connectome–Based Predictive Modeling in Autism | Review Article |

| | | |
|---|---|---|
| 2022 | Individual differences in (dis)honesty are represented in the brain's functional connectivity at rest | Not Whole Brain / ROI Level |
| 2022 | Intrinsic functional connectivity in the default mode network predicts mnemonic discrimination: A connectome-based modeling approach | Not Whole Brain / ROI Level |
| 2022 | Resting-state functional connectivity of social brain regions predicts motivated dishonesty | Not Whole Brain / ROI Level |
| 2023 | A machine learning based approach towards high-dimensional mediation analysis | Substantially modified CPM |
| 2023 | Bidirectional connectivity alterations in schizophrenia: a multivariate, machine-learning approach | Substantially modified CPM |
| 2023 | Brain networks for temporal adaptation, anticipation, and sensory-motor integration in rhythmic human behavior | CPM on task data |
| 2023 | Connectome-based predictive modeling of empathy in adolescents with and without the low-prosocial emotion specifier | Incompatible Accuracy Reporting |
| 2023 | Connectome-based predictive modeling: A new approach of predicting individual critical thinking ability | Not Whole Brain / ROI Level |
| 2023 | Connectome-based predictive modelling of cognitive reserve using task-based functional connectivity | CPM on task data |
| 2023 | Investigating cognitive neuroscience theories of human intelligence: A connectome-based predictive modeling approach | Not Whole Brain / ROI Level |
| 2023 | Seeing the future: Connectome strength and network efficiency in visual network predict individual ability of episodic future thinking | Not Whole Brain / ROI Level |
| 2024 | Brain-based graph-theoretical predictive modeling to map the trajectory of anhedonia, impulsivity, and hypomania from the human functional connectome | CPM on task data |

**Supplementary Table 2 |** Studies selected for literature review of CPM on resting-state fMRI that were excluded from our comparison, listed with the reason for exclusion. The 31 papers are ordered alphabetically, by year.

### Supplementary Figures

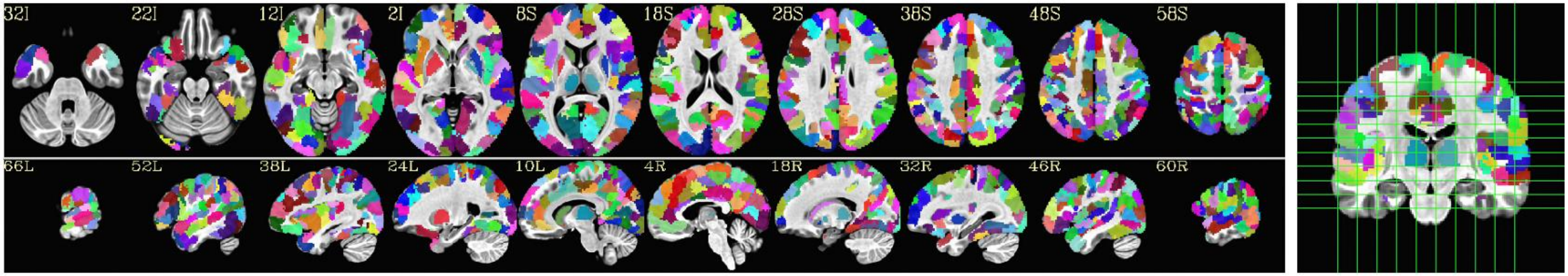


**Supplementary Figure 1 |** Parcellation used to generate FC matrices. This parcellation is a combination of the 400 ROI Schaeffer Atlas plus eight subcortical regions from the AAL atlas. A few ventral regions with poor coverage in the functional scans were removed. The final number of ROIs in the parcellation is 380 regions.

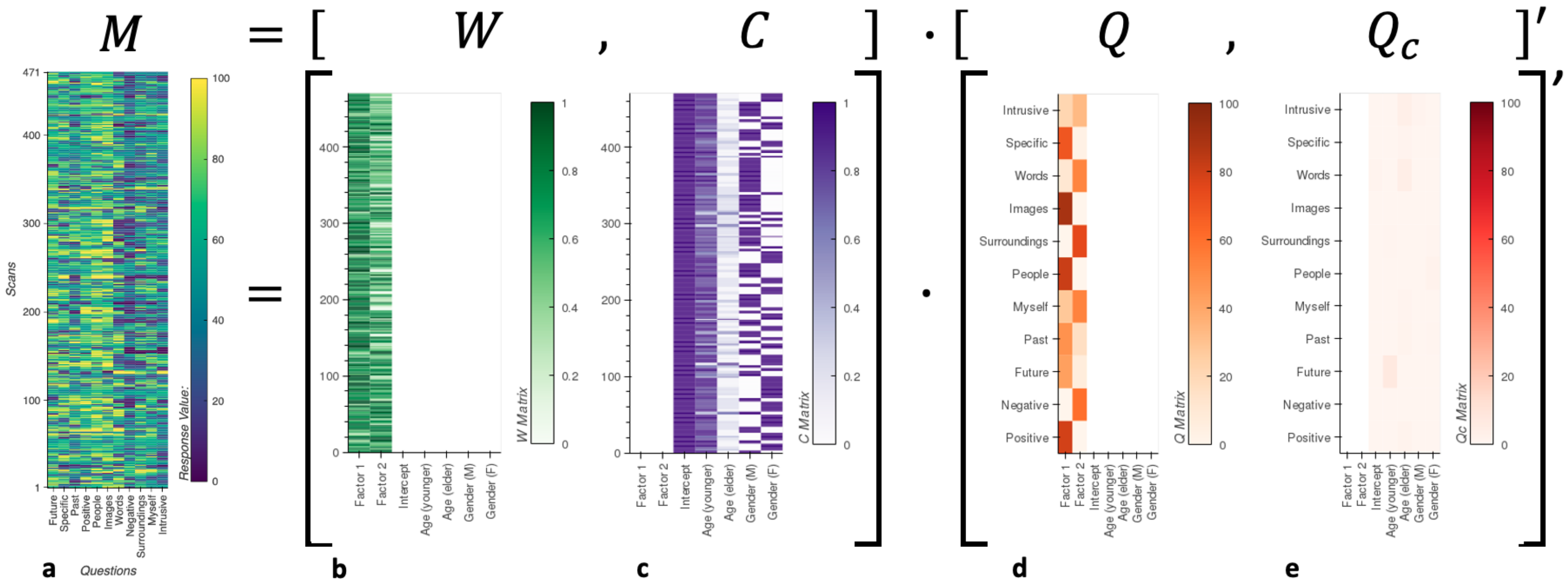


**Supplementary Figure 2 |** Matrix decomposition of questionnaire items using sparse box-constrained non-negative matrix factorization. **a,** matrix M with answers to all 11 questions for all 471 scans. **b,** Matrix W with 2D representation of the questionnaire data. **c,** Matrix C which encoded demographic information (age range and gender). **d,** Matrix Q, which encodes how each of the two dimensions in W (Factor 1 and Factor 2) relate to the original 11 questionnaire items. **e,** Matrix Qc, which contains information about relationships between demographic information and the way participants answered questionnaire items. All entries in the Qc matrix are small compared to those in the Q matrix, which indicate that demographics does not play an important role as a confound in the data.

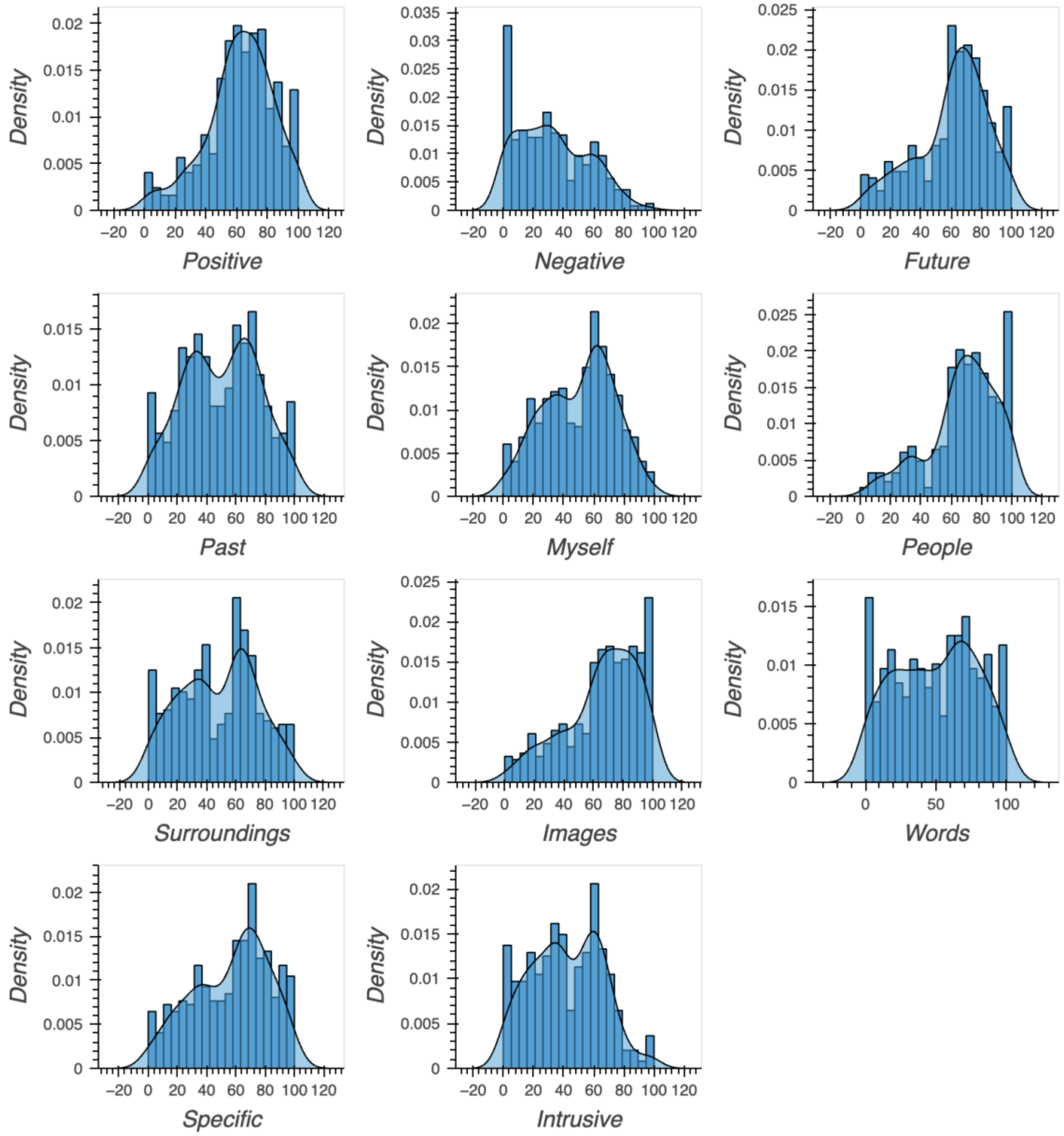


**Supplementary Figure 3 |** Distribution of answers to the 11 questionnaire items about the form and content of thoughts. Labels on the X-axis match those in Table 1 where we provide the original questionnaire items presented to the participants at the conclusion of each resting-state scan.

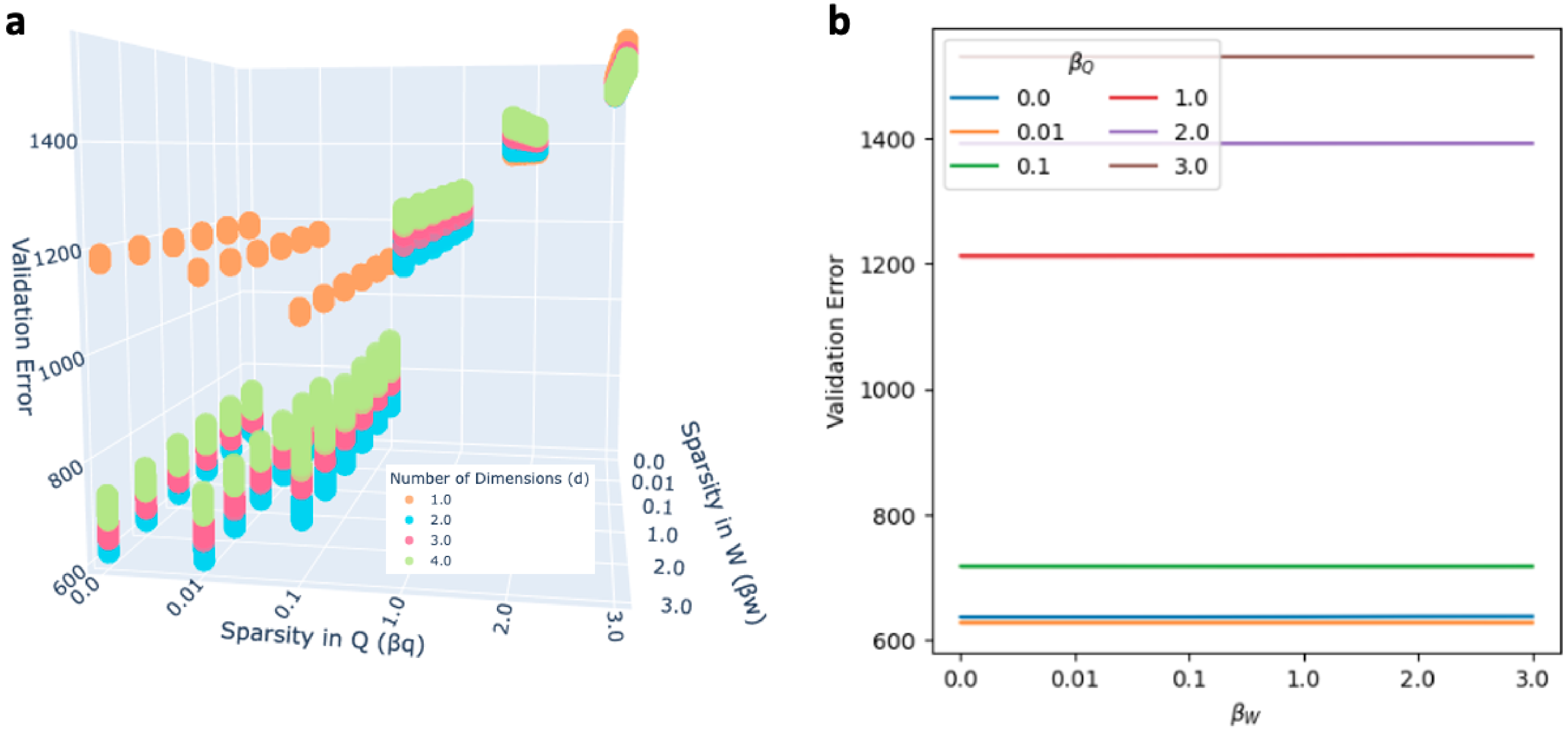


**Supplementary Figure 4 | a,** Validation error cost function for all explored hyper-parameters. Independently of $\beta_Q$ and $\beta_W$, lowest validator errors are always obtained for *d=2*. **b,** Same validation curve errors, but only for d=2. Here we can observe that the lowest error are obtained for $\beta_Q$=0.01 (orange trace). Once $\beta_Q$ is set that way, there is no difference in error across $\beta_W$ values. We selected $\beta_W$ = 0 to avoid unnecessary application of an additional sparsity argument.

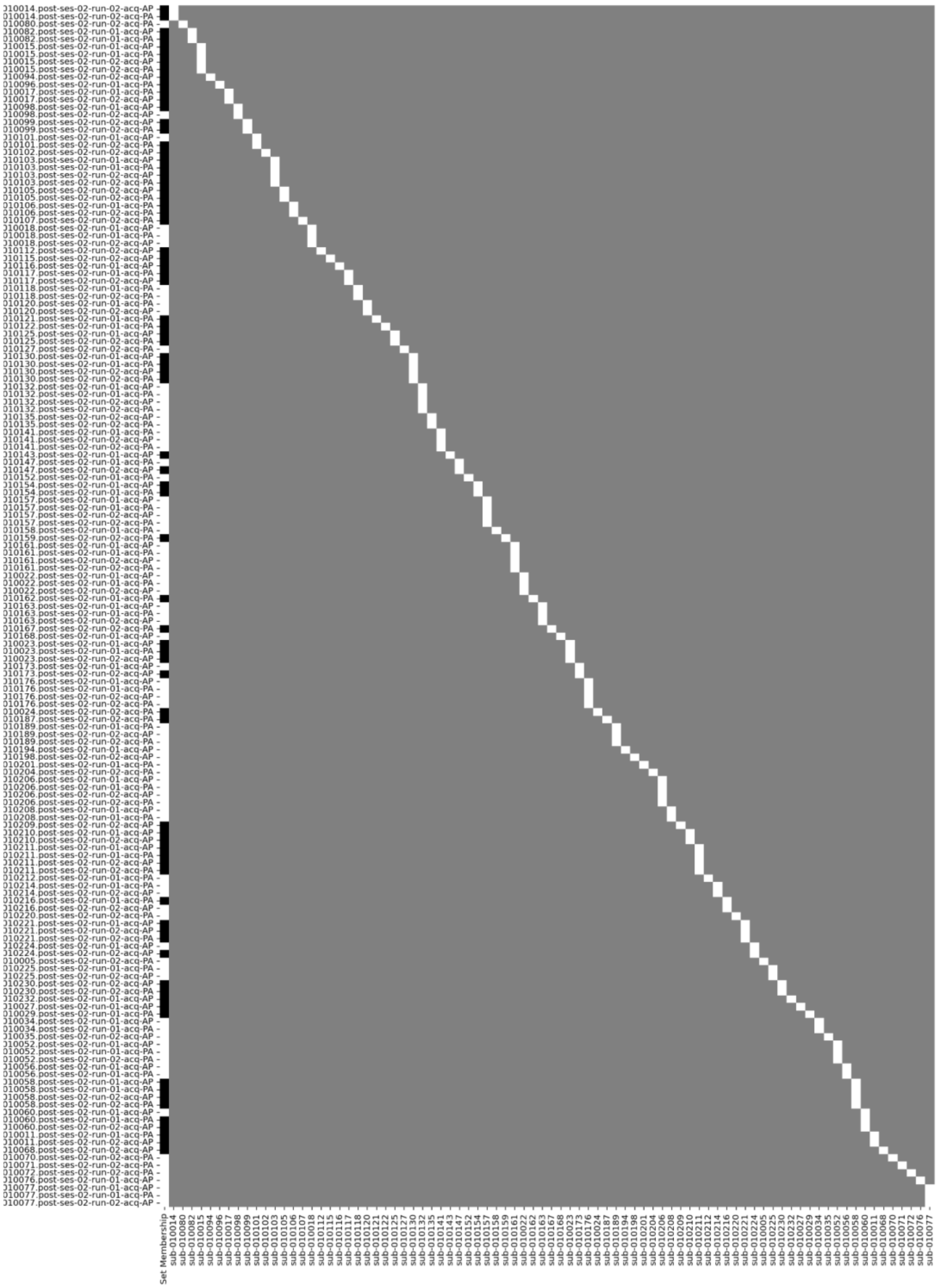

**Supplementary Figure 5 |** Design Matrix for NBS contrast. Rows corresponds to individual scan. First column contains information about set identity (based on TP1 and TP2). The remaining columns encode subject identity for each individual scan.
